# Modeling cardiac fibroblast heterogeneity from human pluripotent stem cell-derived epicardial cells

**DOI:** 10.1101/2023.10.22.563460

**Authors:** Ian Fernandes, Shunsuke Funakoshi, Homaira Hamidzada, Slava Epelman, Gordon Keller

## Abstract

**Abstract (Summary)**

Cardiac fibroblasts play an essential role in the development of the heart and have been implicated in disease progression in the context of fibrosis and regeneration. Here, we established a simple organoid culture platform using human pluripotent stem cell-derived epicardial cells and ventricular cardiomyocytes to study cardiac fibroblasts’ development, maturation and heterogeneity under normal conditions and following treatment with pathological stimuli. We demonstrated that this system models the early interactions between epicardial cells and cardiomyocytes to generate a population of fibroblasts that recapitulates many aspects of fibroblast behaviour *in vivo* including changes associated with maturation and in response to pathological stimuli associated with cardiac injury. Using single cell transcriptomics, we show that the hPSC-derived organoid fibroblast population displays a high degree of heterogeneity that approximates the heterogeneity of populations in both the normal and diseased human heart. Additionally, we identified a unique subpopulation of fibroblasts possessing reparative features previously characterized in the hearts of model organisms. Taken together, our system recapitulates many aspects of human cardiac fibroblast specification, development and maturation providing a platform to investigate the role of these cells in human cardiovascular development and disease.

## Introduction

Cardiac fibroblasts make up approximately 15-30% of the total cells in the heart and have been shown to play important roles in the development of the fetal heart and in maintaining homeostasis in the adult organ^1^. The majority of the cardiac fibroblast population derives from the epicardium, a protective mesothelial layer that surrounds the heart early in development and persists throughout adult life^2^. The epicardium develops from progenitors known as pro-epicardial cells that are specified from a structure positioned near the posterior region of the inferior pole of the developing heart known as the septum transversum mesenchyme^3^. Fibroblasts are generated from the epicardium through an epithelial-to-mesenchymal transition (EMT) that is mediated in part by retinoic acid (RA) secreted from the epicardium and TGFβ produced by the adjacent cardiomyocytes^4^, ^5^. As they are specified, the newly formed fibroblasts migrate to and populate the developing ventricular and atrial myocardium. The cardiac epicardial cells and derivative fibroblasts are distinguished from mesothelial cells and fibroblasts of other organs by the expression of a cohort of genes typically associated with the cardiomyocyte lineage including *GATA4*, *GATA6*, and *HAND2*^6^.

One of the functions of cardiac fibroblasts is the secretion of extracellular matrix proteins that provide support for cardiomyocyte proliferation during fetal life, for the regulation of organ maturation in postnatal life and for structure in the adult heart. Analyses of the matrix at different stages have shown that the composition changes between fetal and adult life as the heart transitions from a developing tissue consisting of proliferative cells to a postnatal organ made up of quiescent cardiomyocytes that have exited cell cycle^7^. During fetal life, the matrix is comprised of a mixture of fibronectin, collagens, and various proteoglycans that functions to support the proliferation of fetal cells. Following birth, there is a shift to a collagen-rich matrix that is less supportive of proliferation and has a greater tensile strength to provide structure for the hemodynamics of the adult cardiac cycle^8^. Currently, it is not known if these changes in the matrix composition reflect the maturation of the population of fibroblasts or are mediated by other cells in the cardiac interstitium with the capacity to produce matrix.

In addition to their function in the normal heart, fibroblasts have been shown to play a key role in the response to tissue damage and diseases of the heart. In model organisms such as the zebrafish that possess cardiac regenerative capacity into adulthood, fibroblasts are activated following injury and deposit a unique fibronectin-rich ECM that is conducive to the proliferation of cardiomyocytes and revascularization of the newly formed myocardium^9^, ^10^. Additionally, the activated cardiac fibroblasts are known to secrete soluble factors such as neuregulin1 (NRG1) and heparin-bound epidermal growth factor (HB-EGF) that promote cardiomyocyte proliferation and the anti-inflammatory molecule IL-10 that modulates inflammation in the damaged tissue^10^, ^11^. As this regenerative phase progresses the activated response of the fibroblasts is downregulated by the secretion of Wntless (WLS), a WNT signaling modulator, by the cardiomyocytes^12^. This transient regulation of the fibrotic response is necessary to maintain the balance of injury resolution with cardiac regeneration^10^.

In adult mammals with little cardiac regenerative capacity, ischemic insults such as myocardial infarction (MI) initiate a response in the fibroblast population that results in the formation of a non-contractile scar that salvages the structural integrity of the myocardium. Fibroblasts play an important role in mediating this repair process through the secretion of tissue-modulating factors and the remodelling of ECM components. Activation of fibroblasts appears to be a common response in other forms of cardiomyopathies as well as advanced stages of these diseases are associated with extensive fibrosis in the heart. Fibrosis can exist as interstitial or replacement fibrosis with the latter being associated with cardiomyocyte necrosis. The role of fibroblasts in these different diseases suggests that activation of the fibroblast population is a common response that persists in the absence of a regeneration program.

Although there is a large body of evidence regarding the dynamics of fibroblast activation and proliferation and scar formation in damaged hearts, the cues regulating these responses are poorly understood. Additionally, it is not known if all resident cardiac fibroblasts respond to injury or if the response is mediated by a specific subset of cells^13^. Recent single cell analyses of adult mouse cardiac fibroblasts have shown that the population is heterogeneous and contains subsets with potential reparative features. Farbehi et al. identified a subpopulation of fibroblasts based on expression of the WNT signaling antagonist WIF1 that appears to promote new blood vessel formation and participates in the regulation of the fibrotic response through the expression of WNT ligands and repressors^14^. In a more recent study, Villalba-Ruiz et al., identified a subpopulation of fibroblasts with similar reparative characteristics. These cells, identified by the expression of an enzyme involved in collagen processing, collagen triple helix repeat-containing protein 1 (CTHRC1), were found to be pro-angiogenic and essential for the scar-forming process^15^. The presence of these subpopulations of fibroblasts suggests that there may be a population, even in the adult context, that possesses the capacity for repair. Whether or not comparable populations exist in the human heart remains to be determined.

Our understanding of human cardiovascular development including the specification of specific cell types such as cardiac fibroblasts is limited due to the scarcity of human heart tissue and our ability to maintain and study it *ex vivo*. Given this, many investigators have turned to human pluripotent stem cells (hPSC) as a source of cardiovascular cells and as a model system to study human cardiovascular development and disease. Over the past decade, protocols have been developed that promote the generation of different cardiomyocyte subtypes as well as many non-cardiomyocyte lineages including epicardial cells from hPSCs ^16-18^. In this study, we used the hPSC system to model the interactions of cardiomyocytes with epicardial cells in 3D organoid structures to recapitulate the events that lead to the development and emergence of the human cardiac fibroblast population. We show that these newly generated fibroblasts seed the organoid tissue, mature in these structures and display responses to pathological stimuli that are similar to those observed in fibroblasts in the failing heart. Single cell RNA sequencing analyses revealed a high degree of heterogeneity within the fibroblast population that recapitulates molecular features of the cardiac fibroblast population in the adult ischemic heart *in vivo*. These analyses also identified a subpopulation of human fibroblasts transcriptionally similar to the reparative cells described in the murine heart post-MI. Together, these findings demonstrate the power of using developmental biology approaches to generate hPSC-derived cardiovascular populations that display functional characteristics and disease responses comparable to the corresponding populations found in the adult heart.

## Results

### Generation of cardiac organoids using ventricular cardiomyocytes and epicardium

To model cardiac fibroblast development and function *in vitro*, we generated organoids consisting of hPSC-derived ventricular cardiomyocytes and epicardial cells. This approach was designed to mimic the known interactions between these two cell types in the early heart that result in the induction of EMT within the epicardial population and the subsequent production of derivative cell types including cardiac fibroblasts^19^. For such models to be accurate and predictable, however, it is important to use cell types that most closely correspond to those from the appropriate staged developing heart. We have previously reported on the generation of hPSC-derived ventricular cardiomyocytes and demonstrated that they represent an immature stage of development^17^, ^20^. In an earlier study, we also described a protocol for the derivation of epicardial cells from hPSCs^16^. While this approach did promote the development of cells with epicardial characteristics, the levels of *ALDH1A2*, a key marker of this lineage, were variable within the population. Additionally, the mesoderm origin of the lineage was not investigated in that study.

To improve the generation of RALDH2^+^ (*ALDH1A2)* epicardial lineage cells, we investigated the role of retinoic acid signalling at the mesoderm stage of development as studies in the mouse and chick suggest that the progenitors of the proepicardium are exposed to retinoid signalling *in vivo*^21^. Additionally, the mesoderm we used to generate epicardial cells expresses RALDH2 indicating that it represents a stage requiring active RA signalling (Supplementary Figure 1A). For these analyses, RALDH2 expressing cells were measured and quantified using the Aldefluor assay^22^. Treatment of the developing population between day 4 and 6 with retinol (ROH), the substrate for RA synthesis, did not alter the proportion of ALDH^+^ cells detected at day 8 of differentiation but significantly increased the proportion measured at day 15 (Figure 1A, B, Supplementary Figure 1B). The addition of ROH did not impact the total cell number, suggesting that RA functions to pattern the RALDH2^+^ lineage at the mesoderm stage of development. The addition of ROH also led to an increase in the expression levels of *ALDH1A2, TCF21* and *GATA5* in the day 15 population (Figure 1C). The expression levels of other genes associated with the epicardial lineage including *WT1*, *TBX18, GATA4*, *GATA6* and *HAND2* were not impacted (Supplementary Figure 1C). These findings show that RA signalling at the mesoderm stage of development enhances the generation of a RALDH2^+^ epicardial population.

**Figure 1.**
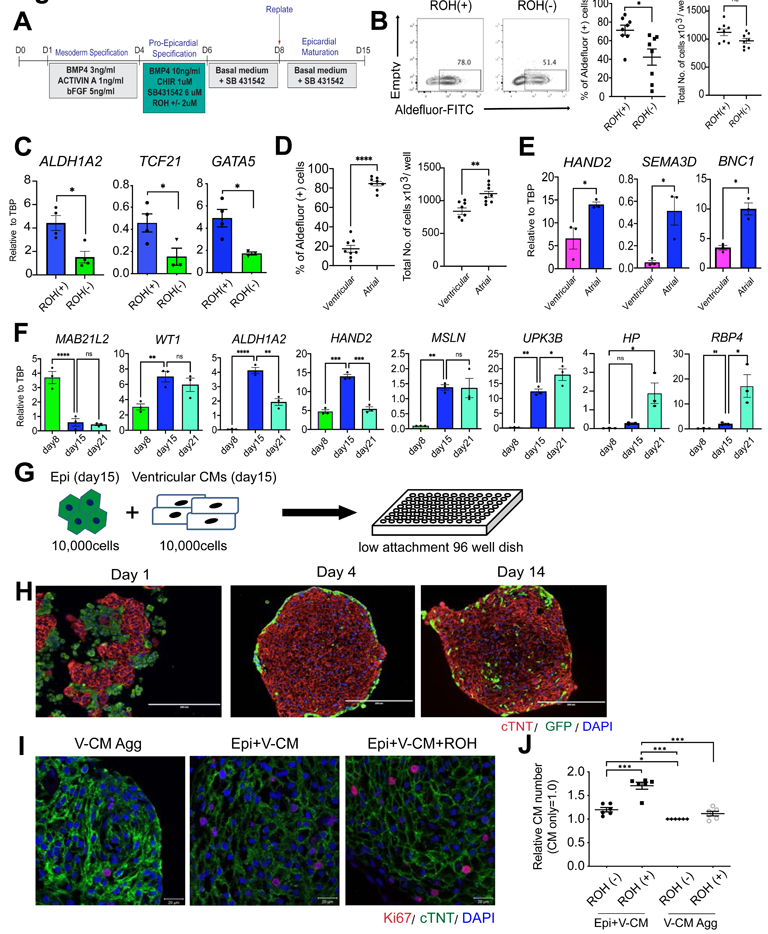
Generation of cardiac organoids with ventricular cardiomyocytes and epicardial cells. (A) Protocol to generate epicardial cells from human pluripotent stem cells. (B) Left; Representative flow cytometry analyses of Aldeflour (ALDH) in day15 retinol (ROH)-treated (ROH(+)) and untreated (ROH(-)) epicardial cells. Right; Quantification of the proportion of ALDH positive cells and total number of cells in ROH-treated and untreated epicardial population (N=8). *p<0.05 by unpaired t test. (C) RT-qPCR expression analyses of *ALDH1A2*, *TCF21,* and *GATA5* in the indicated populations (N=3-4). *p<0.05 by unpaired t test. (D) Quantification of the proportion of ALDH positive cells and total number of cells in ventricular and atrial mesoderm-derived epicardial cells (N=8). **p<0.01, ****p<0.0001 by unpaired t test. (E) RT-qPCR expression analyses of epicardial marker genes in the indicated populations (N=3). *p<0.05, by unpaired t test. (F) RT-qPCR expression analyses of epicardial marker genes in the indicated time points (N=3). *p<0.05, **p<0.01, ***p<0.001, ****p<0.0001 by one-way ANOVA with Tukey’s multiple comparisons. (G) Schema of the coculture of GFP-positive epicardial cells and ventricular cardiomyocyte derived from hPSCs. (H) Representative immunostaining of day1, 4, and 14 cardiac organoids. Scale bar; 200um. (I) Representative immunostaining of Ki67 in ventricular cardiomyocyte aggregates, cardiac organoids, and cardiac organoids treated with ROH. Scale bar; 20um. (J) Quantification of the relative number of cardiomyocytes in the cardiac organoids and cardiomyocyte aggregates (V-CM Agg) treated as indicated (N=6). The number of cardiomyocytes in V-CM only without ROH (ROH(-)) was defined as 1.0 and the relative number of cardiomyocytes was calculated in each condition. *p<0.05, **p<0.01, ***p<0.001 by one-way ANOVA with Tukey’s multiple comparisons. All error bars represent SEM. All values shown for the PCR analyses are relative to the housekeeping gene TBP. V-CM: ventricular cardiomyocytes. Epi: epicardial cells.

The mesoderm used for the above studies has the same ALDH and CD235a/b expression profile and is induced under the same conditions (low concentrations of BMP4 and Activin A) as the mesoderm that gives rise to atrial cardiomyocytes^17^ (Supplementary Figure 1A). To determine if this mesoderm is the best source of these cells, we compared its epicardial potential to that of ALDH^-^/CD235a/b^+^ mesoderm induced with higher concentrations of these pathway agonists that generates ventricular cardiomyocytes (Supplementary Figure 1D). As shown in Figure 1D, the ALDH^+^/CD235a/b^-^ (atrial) mesoderm gave rise to a higher frequency and higher total number of cells than the ALDH^-^/CD235a/b^+^ (ventricular) mesoderm at day 15 of culture. The population derived from the ALDH^+^/CD235a/b^-^ (atrial) mesoderm also expressed significantly higher levels of genes associated with epicardial development including *HAND2*, *SEMA3D* and *BNC1* than the one generated from the ALDH^-^/CD235a/b^+^ (ventricular) mesoderm (Figure 1E). Together, these findings provide strong evidence that the human epicardial lineage develops from ALDH^+^/CD235a/b^-^ ‘atrial’ mesoderm.

In the mouse embryo, the epicardial lineage develops from the pro-epicardial organ that is derived from a transient structure known as the septum transversum mesenchyme (STM) that can be identified by expression of the transcription factor *MAB21L2^23^*. The expression of *MAB21L2* is downregulated with the development of the proepicardial progenitors and the subsequent specification of the epicardial lineage. The establishment of the epicardium proper is defined by the expression of a cohort of genes including cardiac transcription factors (*WT1*, *HAND2*, *BNC1, GATA4/5, TCF21, SEMA3D*) and markers associated with an epithelial identity (*MSLN* and *UPK3B*)^23^. To stage the hPSC-derived lineage, we carried out a kinetic analysis and found that the STM marker, *MAB21L2* was expressed in the population on day 8 and then downregulated at days 15 and 21 (Figure 1F). The levels of some of the epicardial genes including *WT1*, *ALDH1A2*, *HAND2*, *MSLN* and *UPK3B* were low at day 8, showed a significant upregulation by day 15 and then remained constant or declined by day 21. Other genes were expressed at similar levels at all 3 time points or upregulated in the final stages of culture including several adult epicardial markers such as *HP, RBP4, HAS1* and *NFATC1* (Figure 1F, Supplementary Figure 1E). Taken together, these patterns suggest that the transition from the pro-epicardial identity to the epicardial cells that wrap the heart occurs between days 8 and 15 with the transition to a more adult-like phenotype by day 21 of culture. Given that the levels of the fetal epicardial genes, *ALDH1A2* and *HAND2,* were highest on day 15, we chose to use this stage to co-culture with the ventricular cardiomyocytes (Figure 1F).

To generate cardiomyocyte/epicardial organoids, 10^4^ cardiomyocytes and 10^4^ epicardial cells were mixed at a 1:1 ratio in 96 well round bottom plates (Figure 1G). This format yielded single aggregates per well that were maintained in basal medium with no exogenous factors for two weeks. Immunohistochemistry analyses showed that the cells began to form aggregates within 24 hours of culture. At this early stage, there were no signs of organization within the structures. By day 4, however, there was obvious segregation of the cells, as the majority of the epicardial cells (GFP^+^) were now positioned at the periphery of the aggregates, surrounding the inner population of cardiomyocytes. Following an additional 10 days of culture, this segregation was partially lost as an increased number of GFP-positive cells were detected within the aggregates (Figure 1H). Histological analysis showed that GFP-positive epicardial cells exhibited a rounded shape in the early-stage aggregates however, by day 14, they displayed an elongated morphology (Supplementary Figure 1F).

To determine if this simple organoid model can recapitulate key interactions between the epicardium and myocardium in the embryonic heart, we investigated two different aspects of these interactions. The first was the promotion of cardiomyocyte proliferation by the epicardial cells and the second was the induction of EMT in the epicardial cells following contact with cardiomyocytes. Comparison of organoids to cardiomyocyte aggregates (no epicardial cells, hereafter referred to as ventricular cardiomyocyte aggregates, VCM-aggs) revealed that the former had a higher percentage of Ki67-positive cardiomyocytes and a greater number of total cardiomyocytes than the latter (Figure 1I, J, Supplementary Figure 1G) indicating that the presence of epicardial cells does, indeed increase cardiomyocyte proliferation. The addition of ROH to the cultures led to a further increase in the total number of cardiomyocytes in the organoids but not in VCM-aggs (Figure 1J). The addition of ROH however, did not significantly affect the expression level of immature sarcomere-related (*MYH6*), Ca^2+^ handling-related (*ATP2A2*), and ion channel-related (*HCN2, HCN4, CACNA1C, KCNK1*) genes in cardiomyocytes, suggesting this stimulus with or without the organoid environment does not promote the electrophysiological maturation of VCMs (Supplementary Figure 1H).

### Epicardial-cardiomyocyte interactions in the organoids

As a measure of the initiation of EMT and the development of epicardial-derived cell types (EPDCs), we analyzed the organoids at different time points by flow cytometry for the presence of CD90^+^ mesenchymal cells and ALDH^+^ epicardial cells. For these studies, cardiomyocytes were generated from HES2 hPSCs engineered to constitutively express RFP and the epicardial cells from HES2 WT hPSCs. This strategy enabled us to specifically analyze the progeny from the RFP^-^ epicardial cells (Figure 2A). Prior to culture, approximately half of the epicardial cells were ALDH^+^ and of these, 30% expressed low levels of CD90 (CD90^low^) (Figure 2B). A low frequency of CD90^+^ALDH^-^ cells was also detected. By 4 days of culture, the patterns changed as 50% of the cells were CD90^+^ALDH^+^ and more than 20% were CD90^+^ALDH^-^. The increase in the proportion of CD90^+^ cells suggests that EMT has already been initiated. At 14 days of culture, the organoids contained 3 predominant populations: CD90^high^ALDH^-/low^ cells (85%), CD90^-/low^ALDH^high^ cells (7%) and CD90^-^ALDH^-/low^ cells (8%). The large size of the CD90^high^ population at this stage suggests that most of the epicardial cells had undergone EMT and generated mesenchymal progeny. The upregulation of expression of *SNAI1* and *TWIST1* in the CD90^+^ population supports the interpretation that the cells were generated through a EMT process (Supplementary Figure 2A). To further characterize these populations, we isolated them by FACS and analyzed them for expression of genes indicative of fibroblast and smooth muscle cell development and for the persistence of epicardial cells. RT-PCR analyses showed that the CD90^-/low^ALDH^high^ fraction (red square) expressed epicardial-related genes (*ALDH1A2*, *BNC1)*, the CD90^high^ALDH^-/low^ fraction (blue square) expressed genes associated with fibroblast development including *POSTN*, *FN1*, *COL1A1* whereas the CD90^-^ALDH^-/low^ fraction (green square) displayed an expression profile indicative of the presence of coronary smooth muscle lineage cells (*ACTA2*, *MYH11*, and *MLYK)* (Figure 2C). All three populations expressed cardiac transcription factors, including *GATA4*, *GATA6*, *HAND2*, and *TCF21*, indicating that the EPDCs in the organoids maintained cardiac-specific features (Figure 2C). CD31^+^ endothelial cells were not detected in the populations throughout the culture period, suggesting that our day 15 epicardial cells do not possess the potential to differentiate into these cells under these culture conditions (Supplementary Figure 2B).

**Figure 2.**
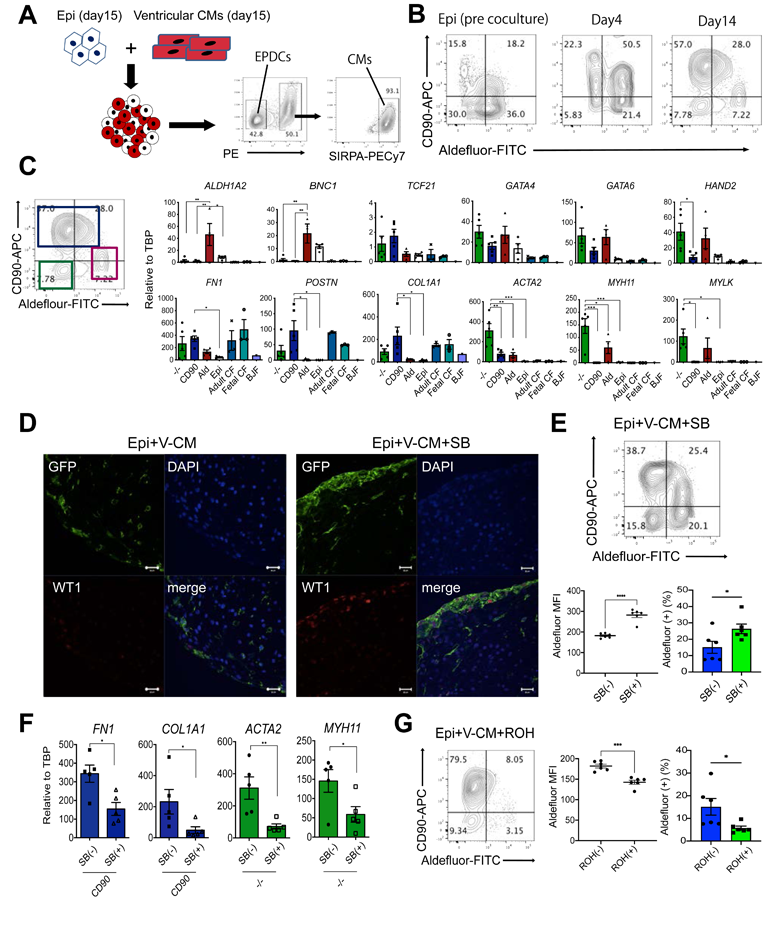
Characterization epicardial-derived cells (EPDCs) in the cardiac organoids. (A) Schema of the generation of the cardiac organoids using the non-labelled epicardial cells and RFP-positive ventricular cardiomyocytes derived from hPSCs, and the strategy to purify the EPDCs and the cardiomyocytes from the cardiac organoids by FACS. (B) Representative flow cytometry analyses of Aldefluor (ALDH) and CD90 in EPDCs in the cardiac organoids in the indicated time points. (C) Left; Representative flow cytometry analyses of 3 subpopulations in EPDCs. Right; RT-qPCR expression analyses of epicardial (*ALDH1A2*, *BNC1*), transcription factor (*TCF21, GATA4, GATA6, HAND2)*, extracellular matrix (*FN1, POSTN, COL1A1*), and smooth muscle (*ACTA2, MYH11, MYLK*) genes in the indicated populations (N=4-5). Adult and fetal cardiac fibroblast (CF) and skin BJ fibroblast were included as a reference. Epi: day15 epicardium prior to coculture, CF: adult and fetal cardiac fibroblasts (N=3 each). BJF: skin BJ fibroblasts (N=1). (D) Representative immunostaining of WT1 and GFP (EPDCs) in the untreated and SB431542 (SB)-treated cardiac organoids. Scale bar; 20um. (E) Upper; Representative flow cytometry analyses of ALDH and CD90 in EPDCs in the SB-treated cardiac organoids. Lower; Quantification of the proportion of ALDH+ cells in each population from the flow cytometry analysis (N=6). (F) RT-qPCR expression analyses of *FN1* and *COL1* in CD90-positive populations in the SB-treated and untreated organoids (N=5) and of *ACTA2* and *MYH11* in ALDH-negative / CD90-negative populations isolated (FACS) from the SB-treated and untreated organoids (N=5). (G) Left; Representative flow cytometry analyses of ALDH and CD90 in EPDCs in the retinol (ROH)-treated cardiac organoids. Right; Quantification of the proportion of each population in the flow cytometry analysis (N=6). *p<0.05, **p<0.01, ***p<0.001, ****p<0.0001 by one-way ANOVA with Tukey’s multiple comparisons in (C) and by unpaired t test in (E)-(G). All error bars represent SEM. V-CM: ventricular cardiomyocytes.

To determine if it was possible to generate fibroblasts with similar characteristics directly from the epicardial cells independent of organoid formation, we treated them with a combination of TGFβ_1_, EGF, bFGF and CHIR for 3 days then with bFGF alone for an additional 3 days (Supplementary Figure 2C). Flow cytometric analyses showed that the majority of the cells within the population following this culture period were CD90^+^ and ALDH^-^, indicative of a transition to a fibroblast fate (Supplementary Figure 2D). RT-qPCR analysis revealed that the cultured population downregulated the epicardial genes *WT1*, *TBX18*, *ALDH1A2* and *BNC1* and upregulated genes associated with fibroblast development including *FN1*, *POSTN* and *COL1A1* to levels similar to those observed in fetal and adult primary cardiac fibroblasts (Supplementary Figure 2E). The transcription factors *GATA4* and *GATA6* were also expressed in the hPSC derived fibroblasts indicating that they displayed the molecular identity of cardiac fibroblasts (Figure 2C, Supplementary 2E).

Immunostaining analyses showed that the organoids but not the VCM-aggs contained fibronectin (Supplementary Figure 2F) indicative of the presence of functional fibroblasts in these structures. MYH11^+^ cells were also found in the organoids, confirming the finding from the RT-PCR analyses that cells with smooth muscle characteristics are generated (Supplementary Figure 2G). Taken together, these observations show that, in the context of the organoids, the epicardial cells undergo EMT and give rise to progeny that display characteristics of cardiac fibroblasts and smooth muscle cells, recapitulating the developmental events observed in the embryonic heart.

Studies in mice have shown that TGFβ signalling plays a pivotal role in the EMT process and in the generation of EPDCs in the developing heart^24^. To determine if this pathway is also involved in these processes in the hPSC-derived organoids, we treated them with the TGFβ inhibitor, SB431542 (SB) and then analyzed the organoids for the presence of epicardial cells and EPDCs. Treatment with SB for 2 weeks resulted in the maintenance of a distinct GFP-positive WT1^+^ epicardial population surrounding the organoid (Figure 2D, Supplementary Figure 2H). Flow cytometric analyses showed that day 14 SB-treated organoids had a higher proportion of ALDH^+^ cells and a lower proportion of CD90^+^ cells than the untreated controls, confirming the presence of a larger epicardial population (Figure 2E vs 2B). RT-PCR analyses of isolated populations revealed that expression levels of *FN1* and *COL1A1* were lower in the SB-treated than in the control CD90^+^ALDH^-^ fibroblasts (Figure 2F). Similarly, the SB-treated CD90^-^ALDH^-^ cells expressed lower levels of *ACTA2* and *MYH11* than the untreated cells. In addition to the TGFΒ pathway, retinoid signalling also appears to promote EMT as organoids treated with ROH were found to contain significantly fewer ALDH^+^ cells than the untreated structures (Figure 2G). RT-PCR analyses of isolated populations showed that expression levels of *FN1*, *COL1A1*, *ACTA2,* and *MYH11* were not significantly different between treated and untreated populations, indicating that retinoid signalling has limited impact on the molecular characteristics of CD90^+^ fibroblast population (Supplementary Figure 2I). Collectively, these observations indicate that EMT in the organoids is regulated in part by TGFβ and retinoid signalling (Supplementary Figure 2J).

### Metabolic maturation of the cardiac organoids

For the organoids to be useful to model diseases that affect the adult heart, the cells should represent a stage of maturation similar to their counterparts in the postnatal organ. As the cells in the organoids and cardiomyocyte aggregates at this stage are immature, we next cultured them in the presence of a PPARα agonist (GW7475), T3 hormone, dexamethasone, and palmitate (PPDT) in low glucose-containing media for an additional 2 weeks, conditions we have previously shown promote metabolic maturation of hPSC-derived cardiomyocytes (Figure 3A)^20^. Flow cytometric analyses of the organoids following this culture period showed that proportion of total RFP^+^ cells (∼50%) and cTNT^+^ MLC2v^+^ cells (85%-90%) within the RFP^+^ subpopulation were similar to that of the 2 week-old organoids (Figure 2A, Supplementary Figure 3A). The cardiomyocyte population in the treated organoid displayed expected characteristics of maturation observed *in vivo* including an increase in sarcomere length within the cells (Figure 3B, Supplementary Figure 3B), a reduction in the proportion of Ki67^+^ cells indicating an exit from the cell cycle (Figure 3C) and an increase in CX43 protein expression (Figure 3D). To functionally assess changes in the oxidative capacity of these populations, we used the Seahorse XF assay to measure the oxygen consumption rate (OCR) between immature and mature organoids and VCM-aggs. Cardiomyocytes cultured in both formats responded to the maturation stimuli as demonstrated by increases in basal respiration, spare capacity and maximal capacity. However, those cultured in the organoids showed higher levels of all these parameters, indicating that the presence of epicardial cells and derivative cell types enhances the metabolic changes in these cells (Figure 3E). Analyses of the treated organoids showed molecular changes within the cardiomyocyte population indicative of maturation, including upregulation of expression of sarcomere-related (*MYOZ2, MYOM3*), Ca^2+^ handling-related (*ATP2A2, HRC, CACNA1C*), mitochondrial (*CKMT2, COX7A1, MFN2*), fatty acid oxidation-related (*MLYCD, CD36, FABP3, CPT1B)* and ion channel-related (*KCNK1, HCN2, KCND3, CACNA1C, KCNJ2, KCNH2*) genes compared to the untreated, age-matched control organoids (Supplementary Figure 3C). The expression levels of immature sarcomere genes (*MYH6* and *TNNI1*) and a pacemaker related ion channel (*HCN4*) were significantly downregulated. However, the expression of adult sarcomere genes (*MYH7* and *TNNI3*) was not significantly different than their immature counterparts suggesting other cues may be missing from the culture to promote these changes.

**Figure 3.**
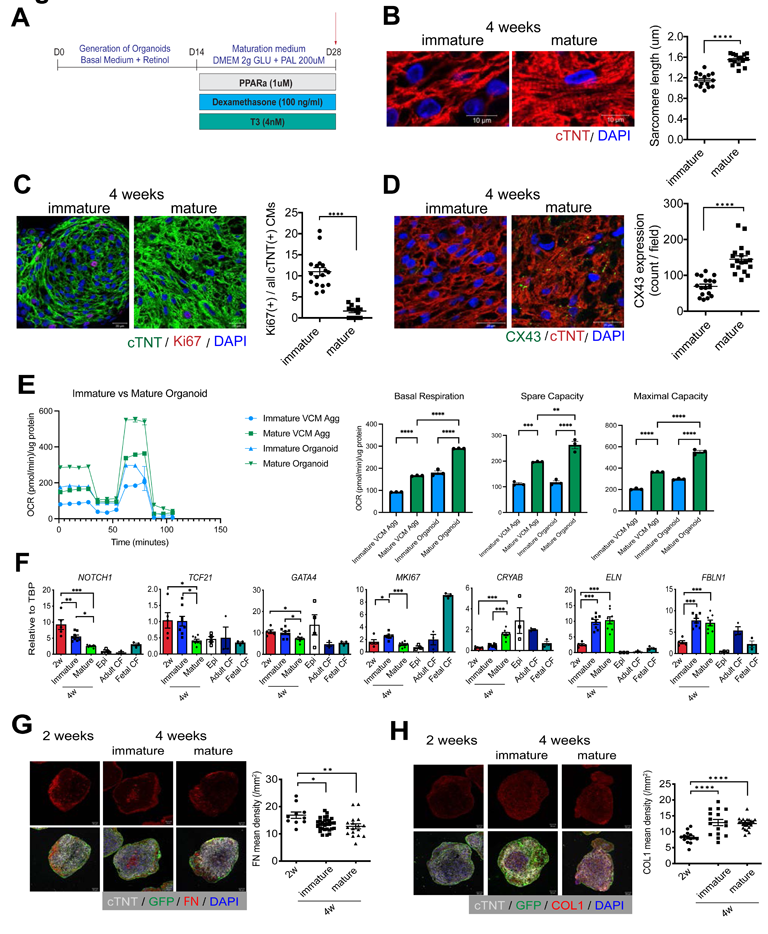
Metabolic maturation of cardiac organoids. (A) Protocol to induce metabolic maturation in the cardiac organoids. (B) Left; Representative immunostaining of cTNT in the immature and mature cardiac organoids. Scale bar; 10um. Right; Quantification of the sarcomere length in the indicated conditions (N=15). (C) Left; Representative immunostaining of Ki67 and cTNT in the immature and mature cardiac organoids. Scale bar; 20um. Right; Quantification of the percentage of Ki67-positive cardiomyocytes in the organoid (N=14-17). (D) Left; Representative immunostaining of CX43 and cTNT in the immature and mature cardiac organoids. Scale bar; 20um. Right; Quantification of the CX43 expression in the organoid (N=17-18). (E) Analyses of the oxygen consumption rate (OCR) in the indicated populations using the Seahorse fatty acid oxidation (FAO) assay (N=3 each). (F) RT-qPCR expression analyses of indicated genes in the early (2 weeks) and late (4 weeks) CD90-positive cardiac fibroblast populations isolated (FACS) from immature and mature organoids (N=5-8). Adult and fetal CF was included as a reference (N=3 each). epi: day15 epicardial cells prior to the initiation of coculture (N=3, 4). (G) Left; Representative immunostaining of fibronectin (FN) in the cardiac organoids at the indicated time points. Scale bar; 20um. Right; Quantification of the FN density at the indicated time points (N=10-30). (H) Left; Representative immunostaining of collagen type 1 (COL1) in the cardiac organoids at the indicated time points. Scale bar; 20um. Right; Quantification of the COL1 density at the indicated time points (N=14-22). *p<0.05, **p<0.01, ***p<0.001, ****p<0.0001 by one-way ANOVA with Tukey’s multiple comparisons in (E)-(H) and by unpaired t test in (B)-(D). All error bars represent SEM. VCM Agg: ventricular cardiomyocyte aggregates. CF: cardiac fibroblasts.

Although the expression levels of genes associated with glycolysis (*GLUT1, GLUT4, PDK4*), were elevated in some experiments, the differences were not consistent and consequently not significant. Together, these findings show that the cardiomyocytes in the organoids underwent changes indicative of metabolic, electrophysiological, and structural maturation, similar to the changes reported in our previous study^20^.

To determine if the changes in culture conditions also promoted maturational changes in the epicardial-derived fibroblast populations, we isolated the CD90^+^ cells from the treated organoids and analyzed them for expression levels of genes that distinguish fetal and postnatal cardiac fibroblast populations^25^, ^26^. This analysis revealed both time-dependent and factor-dependent maturation changes within these cells. Changes independent of the maturation cues included the upregulation of expression of *ELN* and *FBLN1,* genes that encode ECM proteins and the downregulation of *SOX4* and *HBEGF* genes associated with a fetal cardiac fibroblast identity (Figure 3F, Supplementary Figure 3D). These changes are consistent with the changes observed between the fetal and postnatal heart *in vivo*. The expression levels of other genes, such as *NOTCH1, TCF21, GATA4* and *MKI67,* were downregulated following culture in the maturation media, whereas *CRYAB* expression was upregulated (Figure 3F). These differences are also reflective of changes in the fibroblasts in fetal and neonatal hearts. These observations show that the fibroblasts can respond to the same hormonal and metabolic cues governing cardiomyocyte maturation and undergo molecular changes associated with maturation of the population in the developing heart *in vivo*.

To determine if the changes in the fibroblast population were dependent on the environment of the organoid, we subjected the fibroblasts generated directly from epicardial cells to the maturation culture (Supplementary Figure 3E). As these fibroblasts did not survive in the aggregate format, they were cultured and treated in 2D monolayers. RT-qPCR analysis following maturation culture showed that the changes in expression of *NOTCH1, TCF21, GATA4, MKI67* observed in the fibroblasts isolated from the organoid were not detected in the fibroblasts generated directly from the epi cells (Supplementary Figure 3F). The expression of *CRYAB* was significantly upregulated as observed in the organoid fibroblasts and reflective of changes in the fibroblasts in fetal and neonatal hearts. These observations indicate that fibroblasts within the organoids more accurately mimic the *in vivo* maturation process than fibroblasts cultured on their own.

Maturation of the heart is associated with global changes in the ECM composition, characterized by a reduction in fibronectin and an increase in collagen proteins with the transition from fetal to postnatal life^27^. Immunohistochemistry analyses revealed similar changes in the organoids as the fibronectin content decreased between 2 and 4 weeks of culture, whereas the amount of collagen I increased during this timeframe (Figure 3G, H). Culture in the maturation media did not impact these changes.

### Modelling cardiac injury in cardiac organoids

In our previous study, we showed that aggregates of mature cardiomyocytes (without fibroblasts) would undergo changes associated with cardiomyocyte injury following exposure to pathological stimuli including isoproterenol (ISO) and hypoxia^20^. As cardiomyocytes and cardiac fibroblasts are known to interact closely to facilitate remodelling during the progression to heart damage, we next asked if the injury response in the mature organoids more accurately reflected disease progression *in vivo*. In addition to hypoxia and ISO, we also included TGFβ1 as an excess of signalling through this pathway is known to play an important role in the progression of heart failure^28^ (Figure 4A).

**Figure 4.**
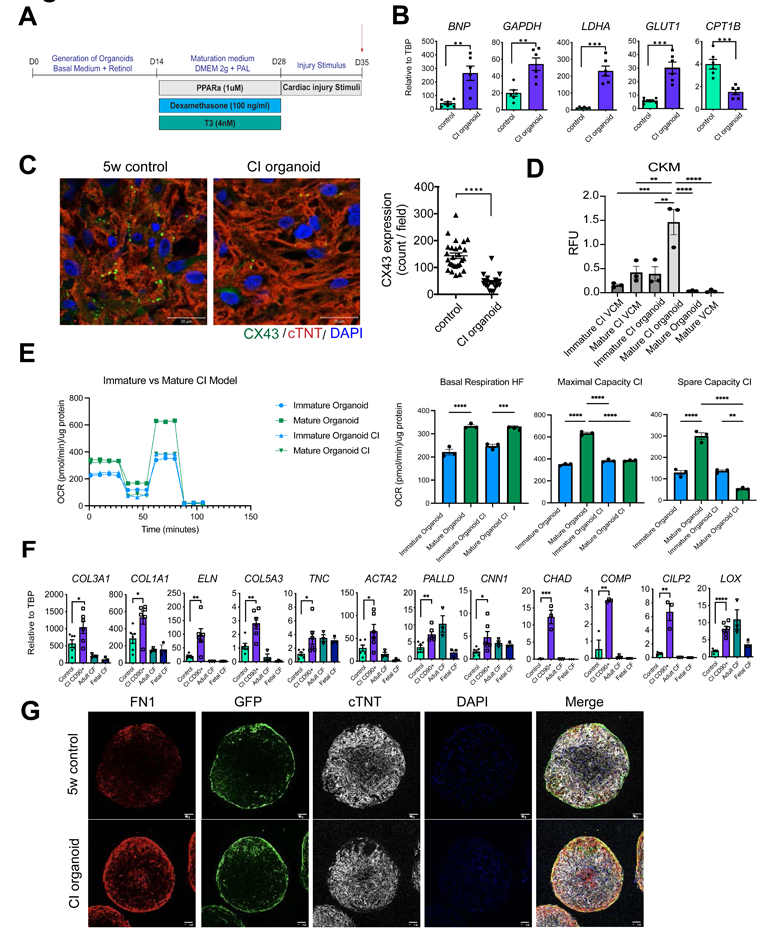
Modeling cardiac injury in cardiac organoids. (A) Protocol to induce cardiac injury in the cardiac organoids. (B) RT-qPCR expression analyses of the heart failure marker (*BNP*) and metabolic-related genes (*GAPDH, LDHA, GLUT1, CPT1B*) in the cardiomyocyte populations isolated from the control (non-treated) and CI organoids (N=6). (C) Left; Representative immunostaining of CX43 in the control and CI-treated organoids. Scale bar; 20um. Right; Quantification of the CX43 expression in the two organoid populations (N=20-27). (D) Quantification of CK release from the indicated CI-treated and control populations (N=3). (E) Analyses of the oxygen consumption rate (OCR) in the indicated populations using the Seahorse fatty acid oxidation (FAO) assay (N=3 each). (F) RT-qPCR expression analyses of extracellular matrix (*COL1A1, COL3A1, ELN, COL5A3*), cytoskeleton (*ACTA2, PALLD, CNN1*), fibrosis-related (*TNC, LOX*), and matrifibrocyte-related (*CHAD, COMP, CILP2*) genes in the FACS isolated CD90^+^ cardiac fibroblast populations from the control and CI organoids (N=3 for matrifibrocyte genes, N=6 for others). Adult and fetal CF RNA was included as a reference (N=3 each). (G) Representative immunostaining of fibronectin (FN) in the control and CI organoids at 5 weeks of culture. Scale bar; 20um. *p<0.05, **p<0.01, ***p<0.001, ****p<0.0001 by one-way ANOVA with Tukey’s multiple comparisons in (D), (E) and by unpaired t test in (B)-(D), (F). All error bars represent SEM. CF: cardiac fibroblasts. CI organoid: cardiac injury organoid. VCM: ventricular cardiomyocytes.

Treatment with ISO/hypoxia/TGFβ_1_ induced an upregulation of the expression of glycolysis-related genes *GAPDH*, *LDHA*, and *GLUT1*, and a reduction in expression of *CPT1B*, a mitochondrial transporter of fatty acids within the cardiomyocyte population (Figure 4B). These changes suggest that the cells are decreasing their capacity for fatty acid oxidation and increasing anaerobic glycolysis, recapitulating the metabolic shift observed in many types of heart disease.

The treated cardiomyocytes also showed elevated levels of *BNP*, a clinical marker of heart failure, a reduction in CX43 expression and downregulation of several calcium handling and ion channel related genes, changes commonly observed in the failing heart ^29^ (Figure 4B, C, Supplementary Figure 4A, B). Notably, the expression level of BNP at baseline in VCM-aggs was relatively higher than that in cardiomyocytes within the organoids and the elevation of BNP by the injury stimuli was not observed in cardiomyocyte aggregates (Supplementary Figure 4C). To further validate the ‘injured’ phenotype of the treated organoid, we next measured CK release, an assay used to detect muscle injury. The outcome of these analyses showed that mature organoids exposed to CI released significantly higher levels of CK than the non-treated mature organoids or the immature organoids. The VCM-aggs did not respond to the CI treatment (Figure 4D). Seahorse analyses showed that the mature organoids treated with CI had significantly reduced maximal and spare OCR capacity compared to the untreated organoids indicating that their capacity for oxidative respiration is reduced, a change observed in cardiomyocytes in the failing heart (Figure 4E). No differences were detected in the treated and non-treated immature organoids. Collectively, these findings show that cells in the mature organoids most accurately recapitulate these key changes associated with cardiovascular disease.

Treatment with the stimuli also had a profound effect on the CD90^+^ fibroblasts as they showed increased expression levels of many ECM genes including *COL1A1, COL3A1*, *ELN* and *COL5A3* (Figure 4F). Many of these genes have been found to be upregulated in the failing heart. The expression levels of genes indicative of activated fibroblasts (myofibroblasts) such as *TNC, ACTA2*, *PALLD*, and *CNN1* as well as those associated with transition to the matrifibrocyte state including *CHAD, COMP,* and *CILP2,* were also upregulated in these fibroblasts^30^. Finally, the expression level of *LOX*, the gene encoding lysyl oxidase, an enzyme inversely correlated with LV ejection fraction in patients with heart failure, was also increased in the activated fibroblasts^31^. Changes in expression of the majority of these fibrosis-related genes were not observed in CI-treated fibroblasts generated directly from the epi cells (Supplementary Figure 4D, E). Immunohistochemical analyses confirmed the gene expression changes in the organoids and showed increased deposition of FN, COL1, and COL3 in the treated organoid compared to the untreated ones (Figure 4G, Supplementary Figure 4F, G). These images clearly demonstrate that the increased FN1 deposition is co-localized with the GFP^+^ population, suggesting that fibroblasts are the main source of ECM production. Collectively, these findings provide evidence that the cardiomyocyte and fibroblast populations within the organoids respond to injury stimuli and faithfully recapitulate changes associated with cardiac injury.

### Single-cell RNA sequencing analyses of the cardiac organoids

To assess the cellular heterogeneity of our organoid populations we performed single-cell RNA sequencing (scRNAseq) at 5 weeks of culture of control and cardiac injury (CI) organoids. To track the RFP labelled cardiomyocyte and GFP labelled epicardial-derived stromal populations, we used the lipid-based multiplexing strategy MULTI-seq, as previously described^32^ (Supplementary Figure 5A). With this approach, we showed that as expected most stromal (cTNT-negative) cells were GFP-positive whereas the cardiomyocytes (cTNT-positive) were RFP-positive (Supplementary Figure 5B). The RFP-positive population from the cardiomyocyte fraction contained some contaminating fibroblasts, which were excluded from downstream analyses. Clustering analysis identified seven cell types and two proliferating populations (Figure 5A, B, Supplementary Table 1). Analyses of the epicardial-derived lineages revealed that both the control and CI organoid contained clusters with expression profiles indicative of pericytes/smooth muscle cells, including elevated levels of *ACTN1, CALD1, PDGFRB, RGS5*, and *TPM2* (Supplementary Figure 5C). The finding that the majority of the EPDC population was THY1 (CD90)-positive fibroblasts corroborated our flow cytometry results. These cells broadly expressed ECM genes, such as *COL1A1, COL3A1, FN1* and *POSTN* indicative of the fibroblast lineage (Figure 2B, C, Supplementary Figure 5C). We additionally identified a subpopulation (cluster 6) that expressed several adipocyte markers including *PPARG, KLF5, BMP2, CEBPA, CD24*, and *DGAT2* confirming the that epicardial derivatives in our cardiac organoids contained adipocyte-like cells in addition to fibroblasts and smooth muscle cells, recapitulating the developmental potential of the epicardium *in vivo*^33^ (Supplementary Figure 5D).

**Figure 5.**
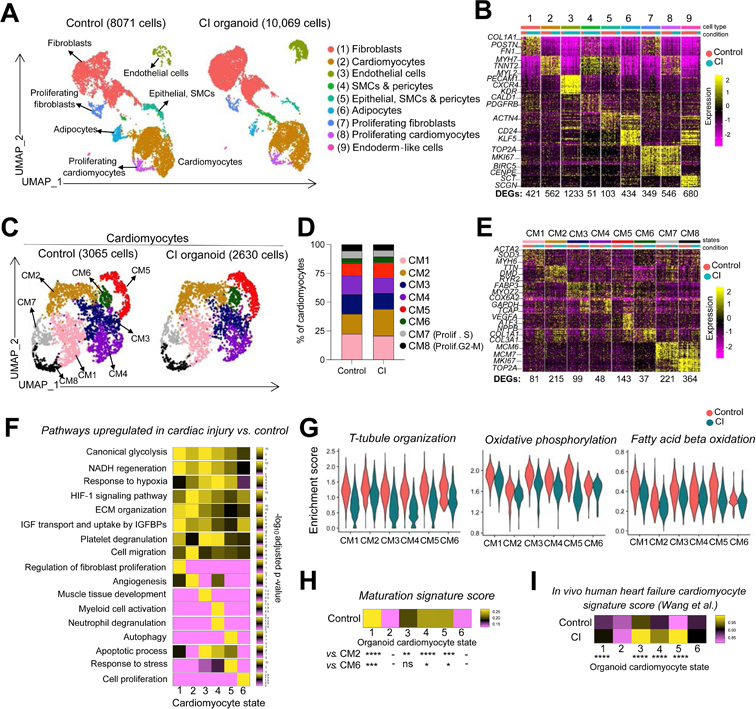
Single-cell RNA sequencing analyses of the cardiac organoids. (A) UMAP dimensionality reduction of total cells in control and cardiac injury (CI) samples. Clusters were annotated based on canonical cell type markers. (B) Heatmap depicting top 30 genes in each cluster in (A) (logFC threshold = 0.3, min.pct = 0.3, adjusted p-value <0.05). The number of DEGs in each cluster relative to all other clusters is indicated below the heatmap. Color bars indicate cell type (top) and condition (bottom). (C) Cardiomyocytes were sub-clustered. UMAP dimensionality reduction of cardiomyocytes in control and CI populations. (D) Relative frequency of each cardiomyocyte cluster in control and CI organoid populations. (E) Heatmap depicting top 30 genes in each cardiomyocyte cluster (logFC threshold = 0.3, min.pct = 0.3, adjusted p-value <0.05). The number of DEGs in each cluster relative to all other clusters is indicated below the heatmap. Color bars indicate cluster(top) and condition (bottom). (F) Pathway enrichment analysis (gProfiler, GO: Biological Processes) was performed using DEGs upregulated in cardiac injury vs. control in each cardiomyocyte cluster. Heatmap depicts the enrichment score (-log10 of adjusted p-value) scaled across clusters for each pathway individually. (G) Pathway scores were generated using genes annotated for each term in the GO: Biological Processes database. Violin plots show single cell expression of each cumulative pathway score in each cluster for control and CI populations. (H) A cardiomyocyte maturation score was generated using 8 maturation signature genes (Funakoshi *et al*. 2021). Heatmap displays the maturation signature score in each cardiomyocyte cluster in the control sample. One-way ANOVA was performed for each cluster compared to the lowest scoring cluster (CM2 and CM6). (I) An *in vivo* human heart failure signature was generated using the top 25 DEGs upregulated in ventricular cardiomyocytes in coronary heart failure vs. normal hearts (single cell dataset published in Wang *et al*.). Each cell in all clusters of control and CI organoids were scored using this signature, and displayed in the heatmap. An unpaired t-test was performed for each pairwise comparison (cardiac injury vs. control in each cluster). SMC: smooth muscle cells, CM: cardiomyocyte, CI organoid: cardiac injury organoid, DEG: differentially expressed genes. ****p<0.0001, *** p<0.001, ** p<0.01, *p<0.05.

We first focused on cardiomyocytes and identified eight transcriptionally distinct clusters in both control and CI organoids (Figure 5C, D, Supplementary Table 2). These cardiomyocyte subpopulations were enriched in a variety of distinct pathways, including wound healing, apoptotic process, stress response and angiogenesis (Supplementary Figure 5E). Clusters 7 and 8 represent proliferating cardiomyocytes marked by the expression of *MKI67* and *TOP2A* (Figure 5E).

To identify transcriptional changes and pathways involved in the cardiac injury response, we selected differentially expressed genes in each cluster between control and CI organoids and performed pathway enrichment analysis. This analysis uncovered changes common to all clusters as well as those unique to specific subsets of cells. Changes found in all clusters included an upregulation of genes related to glycolysis, NADH regeneration and response to hypoxia and a downregulation of genes associated with t-tubule organization, oxidative phosphorylation, and fatty acid beta-oxidation (Figure 5F, G). Significant increases in the levels of *NPPA* and *NPPB* expression were also detected in the CI-treated organoids compared to untreated controls (Supplementary Figure 5F). These findings indicate that the cardiomyocytes in all the clusters recapitulate changes commonly observed in heart failure *in vivo* including the reactivation of a fetal gene signature. Changes specific to a select subset of clusters included the upregulation of expression of genes involved in angiogenesis in cluster 2, inflammation in cluster 4, autophagy in cluster 5, and apoptotic processes in clusters 3, 4 and 5 (Figure 5F).

To determine if these responses correlated with the maturation state of the cells within a particular cluster, we developed a metabolic maturation score using the maturation signature identified in our previous study^20^ and applied it to the different clusters in the organoid population. This analysis showed there was a correlation with maturation as the cells in clusters 1, 3, 4, and 5 that showed the above expression patterns contained the most metabolically mature cardiomyocytes (Figure 5H). Integrated analyses between the profiles of the control and CI-treated organoid and the single-cell RNA seq data on cardiomyocytes from healthy and coronary heart failure patients reported by Wang et. al. showed overlap on the UMAP embedding suggesting transcriptional similarities are present (Supplementary Figure 5G) ^33^.

To further compare the molecular changes observed in the CI organoids to those found in cardiomyocytes isolated from failing hearts, we generated an *in vivo* human heart failure signature based on the data set from the study of Wang et al^34^ and scored cardiomyocytes in each of 6 clusters in the organoids. As shown in Figure 5I, the clusters containing the most mature cells (cluster 1, 3, 4, and 5) in the CI-treated organoids showed the highest degree of transcriptional similarity to the heart failure cardiomyocytes. Additionally, the clusters in the CI organoids were more comparable to the primary heart failure cardiomyocytes than those in the control non-treated organoids. These data suggest that responses of the cardiomyocytes to the pathological stimuli in the organoids recapitulate, to some degree, changes associated with the failing adult heart. Collectively, these findings show that it is possible to model key molecular changes associated with heart failure in cardiac organoids *in vitro* and that these changes are most accurately recapitulated in metabolically mature cardiomyocytes.

### Characterization of cardiac fibroblast heterogeneity in cardiac organoids

To establish the organ identity of the fibroblasts generated in our cardiac organoids, we compared the expression profiles of those from the control organoids to the expression profiles of fibroblasts from different human fetal tissues. For these analyses, we used a tissue-specific fibroblast signature score that we developed by using the top 100 DEGs amongst the fibroblasts from each tissue, based on data from the human fetal cell atlas^35^ and compared it to the organoid fibroblasts. Organoid fibroblasts were most similar to those in the heart, indicating that they display transcriptional features of *in vivo* cardiac fibroblasts (Supplementary Figure 6A).

To characterize the transcriptional heterogeneity present in the fibroblast population, we subclustered COL1A1-positive and GFP-positive cells from the control and CI organoid (Figure 6A, Supplementary Table 3). We identified 12 transcriptional fibroblast states and one proliferating population (MKI67-positive) in both populations. The CI organoids showed an increase in frequency in clusters 1, 2, 3, 4, and 5 and a reduction in frequency of clusters 6, 12, and 13 (Figure 6B, C). Pathway enrichment analysis of the clusters in both control and CI populations showed heterogeneous development of fibroblasts with a wide range of biological processes common to both including heart contraction, regulation of apoptotic processes, angiogenesis, and heart development (Supplementary Figure 6B). To compare the expression patterns of the fibroblasts in the control and CI organoids to those of fibroblasts isolated from normal and failing adult human hearts, we integrated the hPSC-derived subpopulations with their *in vivo* counterparts. These analyses showed the transcriptional similarity between fibroblasts within the organoids and those found in the adult human heart (Supplementary Figure 6C).

**Figure 6.**
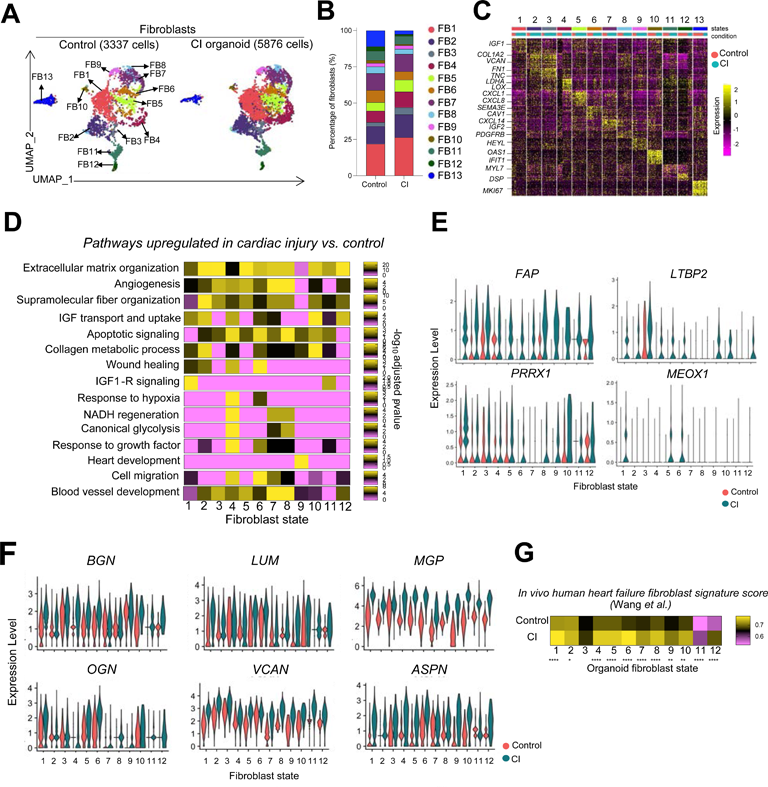
Molecular characterization of fibroblasts in the cardiac organoids. (A) UMAP dimensionality reduction of fibroblasts in control and heart failure samples. (B) Relative frequency of each fibroblast cluster in control and CI samples. (C) Heatmap depicting top 30 genes in each fibroblast cluster (logFC threshold = 0.3, min.pct = 0.3, adjusted p-value <0.05). The number of DEGs in each cluster relative to all other clusters is indicated below the heatmap. Color bars indicate cluster (top) and condition (bottom). (D) Pathway enrichment analysis (gProfiler, GO: Biological Processes) was performed using DEGs upregulated in CI vs. control in each fibroblast cluster. Heatmap depicting the enrichment score (-log10 of adjusted p-value) scaled across clusters for each pathway individually. (E) Violin plots showing expression of fibrosis-related genes (*FAP, LTBP2, PRRX1, MEOX1*) in each fibroblast cluster in the control and CI samples. (F) Violin plots showing expression of proteoglycan-related genes (*BGN, LUM, MGP, PGN, VCAN, ASPN)* in each fibroblast cluster in the control and CI samples. (G) An *in vivo* human heart failure signature was generated using the top 25 DEGs upregulated in cardiac fibroblasts in coronary heart failure vs. normal hearts (single cell dataset published in Wang *et al*.). Each cell in all fibroblast clusters of control and CI organoids were scored using this signature and displayed in the heatmap. An unpaired t-test was performed for each pairwise comparison (cardiac injury *vs*. control in each cluster). CI organoid: cardiac injury organoid.

Pathway analyses of differentially expressed genes between the fibroblast clusters in the control and CI organoids revealed an upregulation of genes involved in ECM organization, collagen metabolic processing, supramolecular fibre organization, angiogenesis, and blood vessel development in most of the clusters in the CI population. Some genes including those associated with wound healing, IGF1-R signaling, response to hypoxia, and cell migration-related genes were upregulated in some specific clusters, suggesting both common and heterogenous responses to the cardiac injury stimuli in the cardiac fibroblast populations (Figure 6D). Analyses of genes known to regulate fibrotic responses such as *FAP, LTBP2, PRXX1* and *MEOX1*^10^, ^36^ those indicative of fibroblast activation, *COL1A1, COL3A1, ACTA2*, and *CNN1,* and those associated with the matrifibrocyte state including *CILP2* and *COMP* were all expressed at higher levels in the fibroblasts from the CI organoids than those from the controls indicating that the stimuli used triggered a fibrotic response (Figure 6E, Supplementary Figure 6D). Additionally, we observed elevated levels of expression of various chondroitin sulphate proteoglycans such as *LUM, BGN, MGP, OGN, VCAN* and *ASPN* in the fibroblasts in the CI organoids (Figure 6F). These changes are in line with findings from a recent study that showed increased levels of these proteoglycans in fibroblasts of the adult human failing heart^37^.

We next compared the molecular profiles of both the control and CI organoids to the top DEGs in the fibroblast cluster from patients with ischemic heart failure to determine if molecular changes in the organoid fibroblasts reflected changes associated with heart disease^34^. As shown in Figure 6G, the profiles from the ischemic heart fibroblasts aligned more closely to the fibroblasts from the CI organoids than to those from the control organoids. Together, these findings demonstrate that the organoid model can recapitulate molecular changes in the fibroblast population associated with ischemic heart disease.

### Identification of a CD9^+^ reparative like fibroblast in the injured heart

To determine if the CI organoids contain the human equivalent of reparative fibroblasts, we generated a molecular score for the CTHRC^+^ reparative fibroblasts (RCF score) using the dataset reported by Ruiz-Villalba et al.^15^ and applied it to the clusters from the organoids. As shown in Figure 7A, clusters 1-6 and cluster 10 showed higher RCF scores than the other clusters, suggesting that they may contain reparative fibroblasts. To further investigate this, we next compared the reparative clusters (1-6, 10) and non-reparative clusters (7-9, 11, 12) to a transcriptional profile of WNTx^+^ reparative fibroblasts characterized by Farbehi et al.^14^. This comparison showed that the putative reparative clusters of fibroblasts from the organoids expressed significantly higher levels of these genes than the non-reparative (canonical) fibroblasts, further supporting the interpretation that the CI organoids contain a subpopulation of reparative fibroblasts (Supplementary Figure 7A). GO pathway analysis revealed elevated expression levels of genes associated with angiogenesis, blood vessel development, collagen organization and fibroblast proliferation in the reparative compared to the canonical fibroblast population, indicating that these cells display the molecular characteristics of the reparative fibroblasts identified in mouse models of heart failure (Figure 7B).

**Figure 7.**
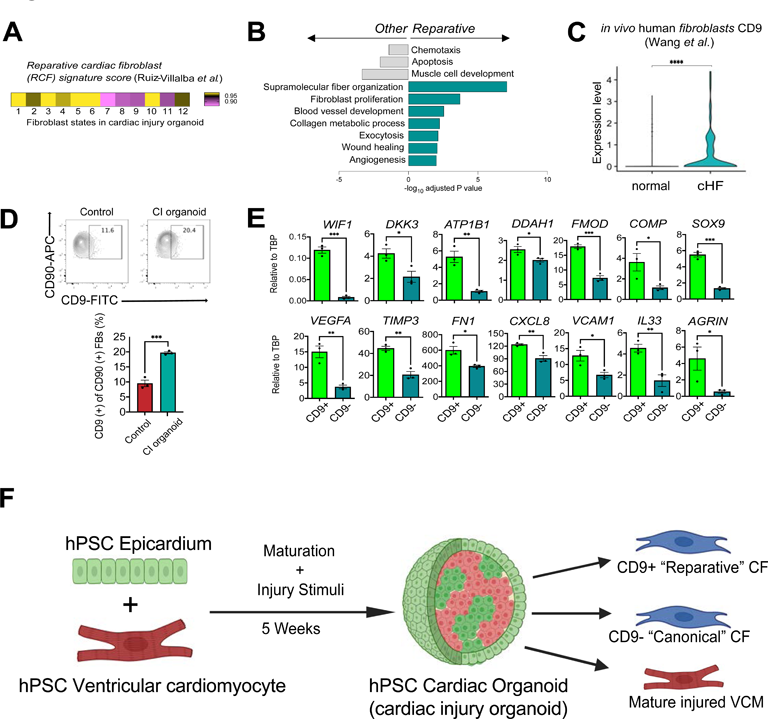
Identification of a CD9^+^ reparative like fibroblast population. (A) A reparative cardiac fibroblast signature was generated using the top 25 DEGs of reparative cardiac fibroblast cluster (as defined by Ruiz-Villalba *et al.*) *vs*. other fibroblasts in Ruiz-Villalba *et al.* Each cardiac fibroblast cluster from the CI population was scored with this signature and displayed in the heatmap. (B) Fibroblast clusters were assigned as “reparative” or “other” based on their relative enrichment of the reparative signature in Figure 7A. DEGs were computed between these two groups in the CI organoids and pathway enrichment analysis was performed (gProfiler, GO: Biological Processes). The enrichment score (-log10 of adjusted p-value) of the top pathways in each group are displayed. (C) Violin plots showing expression of *CD9* in human cardiac fibroblasts in healthy hearts and ischemic failing hearts (coronary HF (cHF)) (single cell dataset published in Wang *et al*.). (D) Upper; Representative flow cytometry analyses of CD9 in the CD90^+^ cardiac fibroblast population in the control and CI organoids. Lower; Quantification of the proportion of CD9^+^ cells in the CD90^+^ cardiac fibroblast population in the control and CI organoids (N=3). (E) RT-qPCR expression analyses of indicated genes in the CD9^+^ and CD9^-^ cardiac fibroblast populations isolated (FACS) from the CI organoids (N=3). (F) Schematic summary of cardiac organoid modeling detailing the heterogeneity of canonical and reparative like fibroblasts observed in the cardiac injury conditions. CF: cardiac fibroblast, VCM: ventricular cardiomyocyte, CI organoid: cardiac injury organoid. ****p<0.0001, *** p<0.001, ** p<0.01, *p<0.05, by unpaired t-test.

We next reanalyzed the published mouse data sets for cell surface markers expressed by the reparative fibroblasts. From this analysis, we found that *Cd9* is expressed in both the *Wif1^+^* and *Cthrc1*^+^ reparative cells (Supplementary Figure 7B, C). To determine if CD9 is also expressed in fibroblasts in human ischemic failing hearts we analyzed the fibroblasts from the Wang et al. report^34^ and found a significant upregulation of CD9 in human ischemic cardiomyopathy fibroblasts compared to control (Figure 7C).

Flow cytometric analyses showed that both the control and CI organoids contained a subpopulation of CD90^+^ cells that expressed low levels of CD9. The frequency of CD9^+^ cells in the CI fibroblast population was approximately double that found in the controls (Figure 7D). RT-PCR analyses of FACS isolated CD9 populations from the CI organoids showed that the expression levels of expression genes associated with the subpopulation of reparative fibroblasts in the mouse including *WIF1, DKK3*, *ATP1B1*, *DDAH1, FMOD, SOX9* and *COMP* were significantly higher in the CD9^+^ than in the CD9^-^ cells (Figure 7E). As expected, the levels of *CD9* were higher in the positive fraction confirming separation based on CD9 expression (Supplementary Figure 7D). The CD9^+^ cells also expressed higher levels of genes associated with angiogenesis including *VEGFA, TIMP3, CXCL8*, *FN1* and *VCAM1,* suggesting a role in supporting vascular development. Differences were also detected in the levels of *IL33*, a cytokine that displays cardioprotective effects on cardiomyocytes and *AGRIN,* an extracellular matrix protein that promotes cardiomyocyte proliferation^9^, ^38^, ^39^ (Figure 7E). Other fibroblast related genes including *WT1, TCF21*, *TBX18* and *COL1A1* were expressed at comparable levels in the CD9^+^ and CD9^-^ subpopulations (Supplementary Figure 7D). Together, these expression patterns support the interpretation that the CD9^+^ fraction of the CD90^+^ population in our organoids represents the human equivalent of the reparative fibroblasts described in the mouse and suggest that they may have roles in matrix production, angiogenesis, and repair of cardiac tissue (Figure 7F).

## Discussion

Cardiac fibroblasts play a central role in heart development, in the maintenance of normal heart function throughout adult life and in the progression of different forms of heart disease. Despite their importance in these processes, our understanding of human cardiac fibroblasts, including their interactions with cardiomyocytes, heterogeneity of the population, the regulation of scar formation and the onset and progression of fibrosis in disease states is limited, largely due to the inaccessibility of human heart tissue. In this study, we have established a simple organoid system with hPSC-derived cells to model key aspects of fibroblast development and maturation and their interactions with cardiomyocytes. With this approach, we were able to document the following characteristics of hPSC-derived fibroblasts that recapitulate known fibroblast behaviour and function in the human heart and in the hearts of model organisms. First, we show that the interaction of epicardial cells and cardiomyocytes leads to the specification of fibroblasts that migrate into and seed the organoid tissue where they mature with time in culture or in response to maturation factors. Second, we demonstrate that the fibroblasts within the organoids can respond to pathological stimuli and undergo changes that reflect changes observed in fibroblasts in the failing heart. Third, our single-cell RNA-seq analyses revealed an unappreciated degree of heterogeneity within the fibroblast population and identified a subpopulation that displays molecular profiles of reparative fibroblasts described in the mouse. Together, these findings highlight the power of the hPSC organoid system to accurately model key aspects of heart development and give rise to cell populations that represent those found in both the fetal and adult organ.

Cardiac fibroblasts are generated from epicardial cells through an EMT process initiated by their interaction with the adjacent cardiomyocytes^5^. Here, we modelled this interaction to generate hPSC-derived cardiac fibroblasts, reasoning that this approach would yield a population that most closely represents the fibroblasts found in the human fetal and adult hearts. A key to the success of this strategy was the generation of an epicardial population that was properly patterned and generated from an appropriate mesoderm population. Previous studies on hPSC-derived epicardial cells have used the expression of WT1 as a defining marker of the lineage^40^, ^41^. However, this transcription factor is expressed in different mesothelial populations that derive from different mesoderm subtypes^42^. We monitored Raldh2 (*ALDH1A2)* expression as an additional marker of the epicardial lineage as it is expressed in the fetal epicardium of the mouse, zebrafish and human^43^ and the function of this enzyme is essential for the ROH-induced proliferation of the cardiomyocytes. Our demonstration that the addition of ROH at the mesoderm stage of development enhances the generation of RALDH2^+^ epicardial cells is in line with observations in the mouse and chicks that the epicardial progenitors are exposed to retinoid signalling *in vivo*^44^. The finding that RALDH2^+^ epicardial cells are preferentially generated from mesoderm that gives rise to atrial cardiomyocytes provides a rationale for optimizing early induction stages for generating cells of this lineage. With access to this optimized population, we were able to recapitulate many aspects of epicardial development and function in the organoid model including the segregation of the cells to the periphery of the tissue, the promotion of proliferation of the adjacent cardiomyocytes and the capacity to undergo TGFβ mediated EMT to generate the spectrum of EPDCs found in the adult heart including fibroblasts, smooth muscle cells and adipocytes. Several recent studies have investigated hPSC-derived epicardial cell-cardiomyocyte interactions *in vitro* either in monolayer or 3D organoid culture formats^45^, ^46^. Consistent with our observations, the findings from this study showed that the epicardial cells can promote proliferation of the cardiomyocytes and can undergo EMT to generate derivative mesenchymal cells. However, these studies did not use well characterized populations, nor did they investigate in detail, the derivative lineages generated in the cultures.

The immature status of hPSC-derived cardiomyocytes has been recognized as a major hurdle in their use to model different forms of heart disease and to develop new cell-based therapies to treat them^47^. To address this issue, different strategies including electro-mechanical stimulation and factor-induced metabolic transitions have been developed to promote the maturation changes known to occur in the cardiomyocyte populations *in vivo*^20^, ^48^. In addition to the cardiomyocytes, there is evidence that the cardiac fibroblasts also undergo maturational changes, including changes in composition of the ECM produced and changes in the expression of genes encoding different transcription factors and signaling molecules^26^. Our findings show that the fibroblasts in the organoids do mature and recapitulate these changes. Notably, some of the changes including the expression levels of specific transcription factors *(TCF21, GATA4* and *MKI67)* and *NOTCH* appear to be induced by the maturation factors that we previously identified promote maturation of cardiomyocytes^20^. These changes occurred in parallel with the expected changes in the cardiomyocyte population, demonstrating that the fibroblasts respond to the same cues that regulate metabolic maturation of cardiomyocytes. The expression levels of other genes, in particular those that encode matrix proteins did not change in response to these factors, but rather changed with time in culture. The pathway(s) regulating these changes are currently not known. Access to organoids that contain matured cardiomyocytes and fibroblasts enables one to model disease processes that approximate those of the adult heart. Indeed, we were able to demonstrate that treatment of the organoids with pathological stimuli induced changes in both the cardiomyocyte and fibroblast lineages associated with cardiac injury *in vivo*. Some of these parameters, including the release of CKM and the reduction in oxidative capacity were only observed in the mature organoids, highlighting the importance of using the appropriate cell types and culture format to model disease. Our single cell analyses revealed a high degree of heterogeneity within the fibroblast and cardiomyocyte populations in both the control and CI organoids, recapitulating the heterogeneity of these populations documented in human hearts and in hearts of model organisms^14^, ^15^, ^49^. Our comprehensive analyses of each of the lineages provides new and important insights into the composition of the populations and their response to pathological stimuli. Notable amongst these was the observation that the organoid-generated cardiomyocytes and fibroblasts showed similar molecular profiles with corresponding cells from the adult heart, suggesting that the strategy of generating cells by recapitulating early developmental events combined with appropriate maturation conditions yields populations appropriate for disease modeling. This notion is supported by the observation that the most mature cardiomyocytes from the CI but not from the control organoids showed striking transcriptional similarities to cardiomyocytes from a failing heart. Additional evidence that this model captures relevant disease-associated changes is the finding that subclusters of fibroblasts from the CI organoids align molecularly with fibroblasts from a failing heart. The identification of a CD9^+^ fibroblast population with a reparative molecular signature and the increase in the size of the population in the CI organoids suggest that the function of these cells is conserved between species and that these cells may play some roles in the disease process in humans. Our finding that *Cd9* is expressed in the *Wif1*^+^ and *Cthrc1^+^*reparative cells in the mouse further validate it as a marker of this subpopulation of fibroblasts and the finding that its expression is upregulated in fibroblasts from the human failing heart adds support to the interpretation that these fibroblasts are involved in heart disease. Reparative fibroblasts were first identified and characterized in the mouse based on the expression of genes that encode factors thought to be important in the repair/regenerative processes in the heart including those that regulate WNT signalling and angiogenesis as well as those reported to be cardioprotective (*IL-33*) or capable of promoting proliferation (*AGRIN*)^9^, ^14^, ^15^, ^38^, ^39^. The expression of genes involved in angiogenesis (*VEGFA, FN1*), and those that control the fibrotic response (*WIF1, DKK3, FMOD*) are overlapping with those of fibroblasts in lower organisms that participate in the regenerative process of the damaged heart^50^. While proposed to be reparative, little is known about the actual function of these cells in the mouse or human as methods to isolate them have not been available to date. Our demonstration that this subpopulation of fibroblasts expresses CD9 and can be isolated by FACS provides the first opportunity to access them and study their function in *in vitro* models of cardiac development and in animal models of heart disease.

In summary, through appropriate modeling of early developmental events, we have established a hPSC-based organoid model that recapitulates cardiac fibroblast development and maturation. Access to distinct subpopulations of fibroblasts, including CD9^+^ reparative cells, in both the control and cardiac injury populations provides an unprecedented opportunity to investigate the role of these different cell types in normal cardiovascular development as well as in heart disease. The findings from such studies have the potential to identify subpopulations of fibroblasts beneficial to heart function as well as those that promote disease progression. The identification of such subpopulations will provide a novel platform for identifying new approaches and ultimately new therapeutics to target specific cell types to manipulate fibrotic responses in heart disease. Additionally, the ability to isolate and co-transplant fibroblast subpopulations such as the CD9^+^ reparative cells with cardiomyocytes in infarcted hearts may uncover regenerative properties that promote remuscularization of scar tissue.

## Methods

### Directed differentiation of hPSCs

For ventricular differentiation, we used a modified version of our embryoid body (EB)-based protocol. hPSC populations (HES2, HES2-GFP, HES2-RFP) were dissociated into single cells (TrypLE, ThermoFisher) and re-aggregated to form EBs in StemPro-34 media (ThermoFisher) containing penicillin/streptomycin (1%, ThermoFisher), L-glutamine (2 mM, ThermoFisher), transferrin (150 mg/ml, ROCHE), ascorbic acid (50 mg/ml, Sigma), and monothioglycerol (50 mg/ml, Sigma), ROCK inhibitor Y-27632 (10 uM, TOCRIS) and rhBMP4 (1 ng/ml, R&D) for 24 h on an orbital shaker (70 rpm). On day 1, the EBs were transferred to mesoderm induction media consisting of StemPro-34 with the above supplements (-ROCK inhibitor Y-27632) and rhBMP4 (8 ng/ml), rhActivinA (12 ng/ml, R&D) and rhbFGF (5 ng/ml, R&D). At day 3, the EBs were harvested, dissociated into single cells (TrypLE), and re-aggregated in cardiac mesoderm specification media consisting of StemPro-34, the Wnt inhibitor IWP2 (1 uM, TOCRIS) and rhVEGF (10 ng/mL, R&D). At day 5, the EBs were transferred to StemPro-34 with rhVEGF (5 ng/ml) for another 5 days and then to StemPro34 for another 5 days. Cultures were incubated in a low oxygen environment (5% CO2, 5% O2, 90% N2) for the first 10 days and a normoxic environment (5% CO2, 20% O2) for the following 5 days.

For cardiomyocyte only maturation cultures, from day 16 to day 18, the EBs were transferred to DMEM high glucose media with XAV (4 uM, TOCRIS) and then transferred to maturation media (DMEM containing low glucose (2 g/L) with Palmitic acid (200 uM, Sigma), Dexamethasone (100 ng/ml, Bioshop), T3 hormone (4 nM, Sigma) and GW7647 (PPARA agonist, 1 uM, Sigma) for the following nine days. Finally, the EBs were cultured in DMEM containing low glucose supplemented with Palmitic acid (200 uM) alone for the following five days (a total of 32 days). Cultures were incubated in a low oxygen environment (5% CO2, 5% O2, 90% N2) for the first ten days and a normoxic environment (5% CO2, 20% O2) for the following 22 days. From day 10 to day 32, the EBs were cultured in polyheme-coated low binding 10 cm culture dishes on an orbital shaker (70 rpm).

For epicardial differentiation, we used different concentrations of rhBMP4 (3 ng/ml) and rhActivinA (1 ng/ml) from day 1 to day 4, followed by Retinol (2 uM, Sigma), rhBMP4 (10ng/ml), SB431542 (6uM, Sigma), CHIR (1uM, Tocris) until day 6. From day 6 to 10 cells are cultured in StemPro34 medium. On day 10 cells are dissociated into single cells (ColB for 1 hour 37 °C followed by TrypLE for 5 minutes at 37 °C) and replated at a density of 250k/mL on 10cm culture dishes in StemPro34 with SB431542 (6uM, Sigma) for 5 days.

For cardiac fibroblast differentiation day 15 epicardial cells were dissociated into single cells (ColB for 1 hour 37 °C followed by TrypLE for 5 minutes at 37 °C) and replated at a density of 500k/mL in 12 well plates for 3 days in bFGF (30ng/mL), TGFB1 (1ng/mL), EGF (5ng/mL), and CHIR (1uM, Tocris) in Fibroblast Medium (25%v/v StemPro34 + 75% v/v IMDM + ITSX 1uL/mL). At 3 days medium was replaced with Fibroblast Medium containing bFGF (30ng/mL) for an additional 3 days in culture. For maturation experiments or CI injury experiments cells grown in maturation medium containing PPDT or the CI injury stimuli cocktail.

### Generation of cardiac organoids

Day 15 hPSC epicardial cells and Day 15 hPSC cardiomyocytes were dissociated into single cells and then mixed in a 1:1 ratio of 20,000 cells in total and spun down in a 96-well polyheme coated round bottom plate for 1 minute at 500 rpm forming organoids in the well. These organoids were cultured for two weeks in StemPro34 medium containing penicillin/streptomycin (1%, ThermoFisher), L-glutamine (2 mM, ThermoFisher), transferrin (150 mg/ml, ROCHE), ascorbic acid (50 mg/ml, Sigma), monothioglycerol (50 mg/ml, Sigma) and retinol (2uM, Sigma).

At two weeks organoids were transferred to 10cm polyheme coated dishes in either to DMEM high glucose (4.5 g/L, ThermoFisher) media containing penicillin/streptomycin (1%, ThermoFisher), L-glutamine (2 mM, ThermoFisher), transferrin (150 mg/mL, ROCHE), ascorbic acid (50 mg/ml, Sigma), monothioglycerol (50 mg/ml, Sigma) for *immature culture conditions* or to *maturation culture conditions* (DMEM containing low glucose (2 g/L) with Palmitic acid (200 uM, Sigma), Dexamethasone (100 ng/ml, Bioshop), T3 hormone (4 nM, Sigma) and GW7647 (PPARA agonist, 1 uM, Sigma)) for the following 10 days and then for another 4 days in DMEM containing low glucose (2 g/L) with Palmitate (200 uM) alone.

### Flow cytometry

The EBs were dissociated by incubation in Collagenase type 2 (0.5 mg/mL, Worthington) in HANKs buffer overnight at room temperature followed by TrypLE for 5 min at 37 °C. Cells were stained for 30 min at 4 °C in FACS buffer consisting of PBS with 5% fetal calf serum (FCS) (Wisent) and 0.02% sodium azide. The following antibodies were used for staining: anti-SIRPa-PeCy7 (Biolegend, 1:1000), anti-CD235a/b-APC (1:100), anti-CD9 (Abcam 1:100), and anti-CD90-APC (BD PharMingen, 1:1000). Aldefluor staining was performed as previously described^17^. Stained cells were analyzed or sorted using the LSR II Flow cytometer (BD PharMingen) or Aria III CFI (BD Biosciences). Data were analyzed using FACS DIVA software (BD) and FlowJo software (Tree Star).

### Immunohistochemistry

The EBs were dissociated as described above and the cells were plated onto 24 well culture dishes pre-coated with matrigel (25% v/v, BD PharMingen). Cells were cultured for 2–3 days and were fixed with 4% PFA in PBS for 15 min at room temperature. Cells were permeabilized and blocked with PBS containing 5% donkey serum, 0.1% TritonX. The following antibodies were used for staining: mouse anti-cardiac isoform of cTNT (ThermoFisher Scientific, 1:200), anti-WT1 (Abcam 1:100), anti-GFP (Rockland 1:200), anti-Ki67 (DAKO 1:200), anti-CX43 (Abcam 1:200), anti-FN (Abcam 1:200), anti-COL1 (Abcam 1:200), anti-COL3 (Abcam 1:200), and anti-MYH11 (Abcam 1:100). For detecting unconjugated primary antibodies, the following secondary antibodies were used: donkey anti-mouse IgG-Alexa488 (ThermoFisher, 1:500), donkey anti-rabbit IgG-Alexa555 (ThermoFisher, 1:500), donkey anti-rabbit IgG-Alexa488 (ThermoFisher, 1:500), donkey anti-mouse IgG-Alexa555 (ThermoFisher, 1:500), donkey anti-mouse IgG-Alexa647 (ThermoFisher, 1:500). Cells were stained with primary antibodies in staining buffer consisting of PBS with 0.1% TritonX, and 5% donkey serum overnight at 4 °C. The stained cells were washed with PBS. The cells were then stained with secondary antibodies in PBS containing 0.1% BSA for 1 h at room temperature followed by DAPI staining. For paraffin sections, tissues were fixed by 4% PFA and embedded. After the deparaffinization and rehydration, heat-induced epitope retrieval was performed followed by immunostaining. Sarcomere length was measured in cTNT+ cardiomyocytes randomly selected from 5 to 10 areas and averaged for each cardiac organoid. The average measurement was obtained by using the distances of 4 consecutive Z lines as shown in Supplementary Figure 3B using Image J. CX43 expression was measured by counting the number of CX43+ staining in one field of view (40x magnification) randomly selected from 5 to 10 areas for each cardiac organoid. FN, COL1 and COL3 expression was measured in each cardiac organoid and normalized by the area (/mm^2^). Stained cells were analyzed using an EVOS Microscope (ThermoFisher), Zeiss LSM700 confocal microscope (Zeiss), and Image J software (NIH).

### Quantitative real-time PCR

Total RNA from samples was isolated using RNAqueous-micro Kit including RNase-free DNase treatment (Invitrogen). Isolated RNA was reverse transcribed into cDNA using oligo (dT) primers and random hexamers and iscript Reverse Transcriptase (ThermoFisher). qRT-PCR was performed on an EP Real-Plex MasterCycler (Eppendorf) using a QuantiFast SYBR Green PCR kit (QIAGEN). The copy number of each gene relative to the house keeping gene TBP is shown. Primer sequences are listed in Supplementary Table 4. Commercial adult cFib, fetal cFib (Lonza) and skin BJ fibroblast (Lonza) were included as a reference.

### CKM release assay

Prespecified treatment groups were collected, and cell lysates were obtained using RIPA buffer, collecting both the lysate and culture supernatant. This samples were processed following manufacturer instructions (ABCAM). CKM phosphorylates creatine to yield creatine phosphate, which dissociates into inosine and inorganic phosphate. In this assay inorganic phosphate reacts with ammonium molybdate to produce phosphomolybdic acid, which is reduced to molybdenum blue. The change in absorbance at 660 nm was monitored with a spectrophotometer (SpectraMAX) and CK activity was expressed as RFU/ug of protein.

### Seahorse XF assay

For the Seahorse XF Mitostress test, several organoids or CM aggregates were plated onto an XFe24 cell culture microplate coated with Cell-Tak at 22.6μg/ml 24 h prior to the assay. 12 h prior to the assay, we replaced the culture media with substrate-limited medium substrate-limited medium: DMEM 2g G + with 1% B27 (-INS), 1% antibiotics, 1% Glutamine, 1% L-Carnitine. 45 min prior to the assay, cells were washed twice with KHB (pH 7.4), and 375 μl/well of Seahorse medium (Modified KHB) was added to the cells and the plate was incubated for 45 min at 37 °C. The assay cartridge was loaded with XF Cell Mito Stress Test compounds (3 μM oligomycin, 4 μM FCCP, 0.5 μM rotenone/0.5 μM antimycin A) 30 minutes prior to the assay. Following calibration, the XF Cell Culture Microplate was immediately inserted into the Seahorse XFe Analyzer and the XF Cell Mito Stress Test was run. After the measurement of OCR, EBs were lysed using RIPA buffer and OCR was normalized to protein content of the corresponding sample lysate using the Bradford assay according to manufacturer’s instructions (ThermoFisher). Data were analyzed using Wave software (Agilent).

### Single-cell RNA sequencing

scRNAseq data generated in this study have been deposited at the GEO database under accession code: GSE221500.

### Library preparation and sequencing

Samples were prepared as outlined in the Chromium Single Cell 3’ Reagents Kits User Guide (v3 Chemistry) and loaded onto the v3 10x Chromium. Sequencing libraries were generated and processed as described by 10x Genomics. Sequencing data (HiSeq2500) was pre-processed using Cell Ranger to create expression matrices. Raw base call (BCL) files were demultiplexed into FASTQ files, and reads were aligned using STAR. Reads were then filtered, followed by barcode and UMI counting to generate the feature-barcode matrices. In order to assign cells back to their sample of origin (RFP vs. GFP), count matrices were generated from the lipid-tagged libraries generated using MULTI-seq. Cells were qualified as either singlets, doublets (antibody signal for more than one hashtag), or negative (insufficient antibody signal for either hashtag). Doublets and negative cells were removed from the analysis to avoid ambiguity associated with RFP vs. GFP designation. All singlets proceeded to subsequent analyses.

### Pre-processing and quality control

The R-based (R 3.6.1) package Seurat v3.1 was utilized for all downstream single cell analyses. Data was filtered by removing genes that were not detected in at least three cells, or cells that expressed less than 200 genes. Candidate doublets or multiplets were excluded by removing cells that expressed >6000 genes. Putative dead or lysed cells were excluded by removing cells in which >40% of transcripts mapped to mitochondrial genes. This permissive cutoff was chosen due to the high level of mitochondrial genes in cardiomyocytes. To address the possibility that dissociation of organoids affected gene expression levels, we removed putative stressed or dying cells which expressed a high percentage of transcripts mapping to dissociation-associated genes (30%) as previously reported. For each of these quality control parameters, we removed outlier cells on the basis of the cell distribution for each measure. Control and cardiac injury datasets were first pre-processed and filtered individually, and then merged for all subsequent analyses using the “merge” function in Seurat.

### Normalization, dimensionality reduction and clustering

Data was normalized using SCTransform in Seurat v3.1. SCTransform removes technical variation while preserving biological variation, and leverages regularized negative binomial regression. This function serves to normalize the data, select highly variable features, and scale the data. The variance-stabilizing transformation (vst) method in SCTransform was used to select the top 3000 highly variable genes. The percentage of mitochondrial transcripts and the number of counts were selected as parameters for regression. Principal component analysis (PCA) was used for dimensionality reduction. The most statistically significant PCs were selected using an elbow plot displaying the standard deviation of each PC. In order to account for technical variation from batch effects between the control and heart failure samples, we performed integration of the two datasets using Harmony. Here, PCA embeddings were adjusted and utilized in graph-based clustering with the FindNeighbors and FindClusters commands. Finally, Uniform Manifold Approximation and Projection (UMAP) was utilized for non-linear dimensionality reduction and projection to a 2D plot for visualization. To compare the heterogeneity present in our organoids to those in human fetal or human adult heart failure, we integrated our datasets using Harmony to remove batch effects while preserving biological variation.

### Differential gene expression and pathway enrichment analysis

Clusters were annotated on the basis of differentially expressed genes (DEGs). DEGs were computed with the FindAllMarkers command using the Wilcoxon Rank Sum Test (min.pct: 0.3; logFC threshold: 0.3; adjusted p-value < 0.05). The top 30 DEGs (based on logFC) were selected for each cluster and displayed on a heatmap, which was downsampled to a maximum of 50 cells per cluster for visualization purposes. Datasets were annotated based on canonical cell type markers in the human heart. In order to define genes that are induced in cardiac injury relative to control organoids, we first used the “subset” function to isolate each cluster separately. We then used the FindMarkers function to compute differentially expressed genes across conditions (cardiac injury vs. control; min.pct: 0.3; logFC threshold: 0.3; adjusted p-value < 0.05). Pathway enrichment analysis was performed either cluster-defining DEGs or DEGs induced in cardiac injury for each cluster. We used gProfiler (https://biit.cs.ut.ee/gprofiler/gost) to measure over-representation of our DEGs against the Gene Ontology (GO): Biological Processes database (http://www.geneontology.org). We displayed the -log10 of the adjusted p-value for each GO term as a measure of the pathway enrichment score.

### Quantification and statistical analysis

All data are represented as mean ± standard error of mean (SEM). Indicated sample sizes (n) represent biological replicates including independent cell culture replicates and individual tissue samples. For single cell data, sample size represents the number of cells analyzed from at least three independent experiments. No statistical method was used to predetermine the samples size. Statistical significance was determined using a student’s t test (unpaired, two-tailed) or one-way ANOVA with Tukey’s multiple comparisons in GraphPad Prism 6 software (GraphPad Software). All statistical parameters are reported in the respective figures and figure legends.

---

**Supplementary Figure 1. Generation of cardiac organoids with ventricular cardiomyocytes and epicardial cells.** (A) Left; Representative flow cytometry analyses of Aldefluor and CD235a/b in day 4 atrial mesoderm. Right; RT-qPCR expression analyses of *ALDH1A2* in atrial and ventricular mesoderm (N=4). (B) Left; Representative flow cytometry analysis of Aldefluor in ROH-treated and untreated epicardial cells at day 8. Right; Quantification of the percentage of Aldefluor-positive cells in the indicated conditions (N=5). (C) RT-qPCR expression analyses of epicardial-related and cardiac transcription factors (*TBX18, GATA4, GATA6, HAND2, WT1*) in the ROH-treated and untreated (ROH (-)) epicardial cells (N=3-4). (D) Representative flow cytometry analyses of Aldefluor and CD235a/b in day 4 ventricular mesoderm. (E) RT-qPCR expression analyses of epicardial-related genes (*BNC1, TBX18, GATA4, SEMA3D, TCF21, HOTAIRM1, HAS1, NFATC1*) in the epicardial cells at the indicated time of culture (N=3). (F) Representative immunostaining of GFP and cTNT in day1, 4, 14 cardiac organoids. White arrows indicate elongated GFP-positive EPDCs. GFP; epi-derived cells (EPDCs), Scale bar; 20um. (G) Quantification of the changes in the percentage of Ki67-positive cardiomyocytes in the cardiac organoids and cardiomyocyte aggregates (VCM Agg) treated as indicated (N=6). (H) RT-qPCR expression analyses of sarcomere-related (*MYH6*), Ca^2+^ handling-related (*ATP2A2*) and ion channel-related genes (*HCN2, HCN4, CACNA1C, KCNK1*) in the indicated conditions (N=3-4). *p<0.05, **p<0.01, ***p<0.001 by unpaired t-test (A) and by one-way ANOVA with Tukey’s multiple comparisons (E) and (G). All error bars represent SEM.

**Supplementary Figure 2. Characterization epicardial-derived cells (EPDCs) in the cardiac organoids.** (A) RT-qPCR expression analyses of EMT-related (*SNAI1*, *TWIST1*) genes in the indicated populations (N=4-5). Adult and fetal CF was included as a reference (N=3 each). Epi: day15 epicardium prior to coculture. (B) Representative flow cytometry analysis of Aldefluor and CD31 in EPDCs in the cardiac organoids in the indicated time points. (C) Protocol to directly differentiate hPSC-derived epicardial cells into cardiac fibroblasts. (D) Representative flow cytometry analysis of CD90 and Aldefluor in directly differentiated CFs derived from epicardial cells. (E) RT-qPCR expression analyses of epicardial (*WT1, TBX18, ALDH1A2*, *BNC1*), transcription factor (*TCF21, GATA4, GATA6, HAND2)*, extracellular matrix (*FN1, POSTN, COL1A1*), and smooth muscle (*ACTA2, MYH11, MYLK*) genes in the indicated populations (N=3). Adult and fetal CF were included as a reference (N=3 each). Epi: day15 epicardium, (F) Representative immunostaining of fibronectin (FN) and GFP (EPDCs) in the cardiac organoids and ventricular cardiomyocyte aggregates (V-CM Agg). Scale bar; 20um. (G) Representative immunostaining of MYH11 in the GFP-positive EPDCs in the cardiac organoids. Scale bar; 20um. (H) Representative immunostaining of day 14 SB-treated cardiac organoids. Scale bar; 200um. (I) RT-qPCR expression analyses of *FN1*, *COL1, ACTA2,* and *MYH11* in CD90-positive populations in the ROH-treated and untreated organoids (N=5). (J) Schematic of the cardiac organoids showing the interaction between the cardiomyocytes and the epicardial cells. ****p<0.0001, *** p<0.001, ** p<0.01, *p<0.05, by one-way ANOVA with Tukey’s multiple comparisons in (A) and by unpaired t test in (E)(I).

**Supplementary Figure 3. Metabolic maturation in cardiac organoids.** (A) Representative flow cytometry analysis of cTNT and MLC2v in RFP^+^ sorted population in the cardiac organoids. (B) The method to measure sarcomere length in an image of cTNT staining. Sarcomere length was measured in an area with a white square by Image J. Scale bar; 10um. (C) RT-qPCR expression analyses of the sarcomere (*MYOZ2, MYOM3, DES, MYH7, MYH6, TNNI1, TNNI3*), Ca handling (*ATP2A2, HRC*), fatty acid oxidation (*CD36, FABP3, CPT1B, MLYCD*) and mitochondria (*CKMT2, COX7A1, MFN2*), ion channel (*KCNK1, HCN2, KCND3, CACNA1C, KCNJ2, KCNH2, HCN4*), glycolysis (*GLUT1, GLUT4, PDK4*) -related genes in FACS isolated cardiomyocyte from the immature and mature cardiac organoids and immature and mature ventricular cardiomyocyte aggregates (VCM Agg) (N=3). (D) RT-qPCR expression analyses of *SOX4* and *HBEGF* in FACS isolated CD90^+^ cardiac fibroblasts populations from the immature and mature cardiac organoids (N=4-8). Adult and fetal CF was included as a reference (N=3 each). epi: day15 epicardial cells prior to the initiation of coculture (N=3, 4), CF: cardiac fibroblasts. (E) Protocol to culture directly differentiate hPSC-derived epicardial cells in the maturation media. (F) RT-qPCR expression analyses of indicated genes in the day23 CFs and day30 CFs cultured in the maturation media (N=3). Adult and fetal CF were included as a reference (N=3). epi: day15 epicardial cells (N=3, 4), *p<0.05, **p<0.01, ***p<0.001, ****p<0.0001 by one-way ANOVA with Tukey’s multiple comparisons in (C) (D) (F). All error bars represent SEM.

**Supplementary Figure 4. Modeling cardiac injury in cardiac organoids.** (A) RT-qPCR expression analyses of *GJA1* in the FACS isolated cardiomyocyte populations from the control and CI organoids (N=6). (B) RT-qPCR expression analyses of Ca^2+^ handling-related (*ATP2A2*) and ion channel-related (*HCN4, HCN2, CACNA1C, KCNK1*) in the FACS isolated cardiomyocyte populations from the control and CI organoids (N=3). (C) RT-qPCR expression analyses of heart failure marker (*BNP*) in the ventricular cardiomyocyte aggregates (V-CM-Agg) cultured as indicated (N=5). (D) Protocol to expose the cardiac injury stimuli in directly differentiated CFs derived from hPSC-epicardial cells. (E) RT-qPCR expression analyses of extracellular matrix (*COL1A1, COL3A1, ELN, COL5A3*), cytoskeleton (*ACTA2, PALLD, CNN1*), fibrosis-related (*TNC, LOX*), and matrifibrocyte-related (*CHAD, COMP, CILP2*) genes in the indicated populations (N=3 for matrifibrocyte genes, N=6 for others). Adult and fetal CF was included as a reference (N=3 each). (F) Left; Representative immunostaining of fibronectin (FN) in the control and CI organoids at 5 weeks of culture. Scale bar; 20um. Right; Quantification of the FN density in the two organoid populations (N=19-22). (G) Upper left; Representative immunostaining of collagen type 1 (COL1) in the control and CI organoids at 5 weeks of culture. Scale bar; 20um. Upper right; Quantification of the COL1 density in the two organoid populations (N=20-21). Lower left; Representative immunostaining of collagen type 3 (COL3) in the control and CI organoids at 5 weeks of culture. Scale bar; 20um. Lower right; Quantification of the COL3 density in two organoid populations (N=20-25). CI organoid: cardiac injury organoid. CF: cardiac fibroblasts. *p<0.05, **p<0.01, ***p<0.001, ****p<0.0001 by unpaired t-test. All error bars represent SEM.

**Supplementary Figure 5. Single-cell RNA sequencing analyses of the cardiac organoids.** (A) Flow chart of single-cell RNA sequencing sample workflow. RFP-positive and GFP-positive fractions of control and CI organoids were sorted and labelled individually using lipid-tagged antibodies, and subsequently pooled for sequencing using the 10x Genomics platform and MULTI-seq. Following sequencing, the data was demultiplexed to assign cells back to RFP or GFP fractions. (B) UMAP dimensionality reduction of total cells in control and CI samples split by RFP and GFP fractions. (C) UMAP plots showing expression of smooth muscle/pericyte and fibroblast-related genes (*RGS5, PDGFRB, ACTN1, CALD1, TPM2, COL1A1, COL3A1, FN1, POSTN, THY1*). (D) UMAP plots showing expression of adipocyte-related genes (*PPARG, KLF5, BMP2, CEBPA, CD24, DGAT2*). (E) Pathway enrichment analysis was performed for each cardiomyocyte cluster in Figure 5 (gProfiler, GO: Biological Processes). The top unique pathways in each cluster are displayed with their enrichment score (-log10 of adjusted p-value). (F) Violin plots showing expression of *NPPA* and *NPPB* in each cardiomyocyte cluster in control and CI samples. (G) Integration analysis of cardiomyocyte population in control and CI organoid samples with cardiomyocytes from *in vivo* human coronary heart failure dataset (Wang *et al*.). Datasets were downsampled to 400 cells each to avoid biases associated with differential cell numbers. CM: cardiomyocyte, CI organoid: cardiac injury organoid.

**Supplementary Figure 6. Molecular characterization of fibroblasts in the cardiac organoids.** (A) Single-cell RNA sequence data of human fetal fibroblasts from 5 tissues (Cao *et al*.) were compared to identify tissue-specific fibroblast signatures. Each tissue-specific fibroblast signature was used to score control organoid fibroblasts. One-way ANOVA was performed for each tissue signature compared to the heart signature. ****p<0.0001. (B) Pathway enrichment analysis was performed for each fibroblast cluster in Figure 6 (gProfiler, GO: Biological Processes). The top unique pathways in each cluster are displayed with their enrichment score (-log10 of adjusted p-value). (C) Integration analysis of fibroblast population in control and CI organoid samples with fibroblasts from *in vivo* human coronary heart failure dataset (Wang *et al*.). Datasets were downsampled to 400 cells each to avoid biases associated with differential cell numbers. (D) Violin plots showing expression of fibrosis-related genes (*COL1A1, COL3A1, ACTA2, CNN1, CILP2, COMP*) in each fibroblast cluster in control and CI samples. FB: fibroblast, CI organoid: cardiac injury organoid.

**Supplementary Figure 7. Identification of a CD9^+^ reparative like fibroblast population** (A) A signature for the WNTx fibroblast cluster defined by Farbehi *et al*. was generated by comparing the WTNx cluster to all other fibroblasts. “Reparative” and “other” fibroblasts from the CI organoids (see Figure 7) were scored with this signature. An unpaired t-test was performed to compare the two groups. (B) Clustering analysis of fibroblast populations and UMAP plots showing expression of *Wif1* and *Cd9* in mouse heart *in vivo* (Farberi *et al*.) (C) Left; Clustering analysis of fibroblast populations and UMAP plots showing expression of *Cthrc1*. Right; Violin plots showing expression of *Cd9* in Cthrc1-positive fibroblasts at the indicated time points. All data are analyzed from the mouse heart dataset of Ruiz Villalba *et al*. (D) RT-qPCR expression analyses of indicated genes in the CD9^+^ and CD9^-^ cardiac fibroblast populations isolated from the CI organoids (N=3). CI organoid: cardiac injury organoid. ****p<0.0001, *** p<0.001,** p<0.01, *p<0.05, by unpaired t-test.

**Supplementary Figure 8. Gating strategy for flow-cytometric analysis.** (A) Gating strategy for sorting RFP(+) / SIRPA(+)-cardiomyocytes and CD90(+)-fibroblasts derived from epicardial cells in the cardiac organoid used in Figure 2-4 and Supplementary Figure 2-4. Representative flow cytometry result of CD90 and Aldefluor in EPDC population in the 4-week cardiac organoids are shown. (B) Representative flow cytometry analysis of CD90 and Aldefluor in EPDC population in the 5-week cardiac organoids with or without cardiac injury. (C) Gating strategy for sorting RFP-positive and GFP-positive fractions of the cardiac organoids for MULTI-seq shown in Figure 5, 6 and Supplementary Figure 5, 6.

## Supplementary Tables

**Supplementary Table 1:** Differentially expressed genes in each cell type in the combined analysis of all cells in control and cardiac injury organoids.

**Supplementary Table 2:** Differentially expressed genes in each cardiomyocyte cluster and genes induced in the cardiac injury versus control organoids.

**Supplementary Table 3:** Differentially expressed genes in each fibroblast cluster and genes induced in cardiac injury versus control organoids.

**Supplementary Table 4:** Primer lists for RT-qPCR.

