## Supplementary Figure for "Modeling cardiac fibroblast heterogeneity from human pluripotent stem cell-derived epicardial cells"

### Supplementary Figure 1.

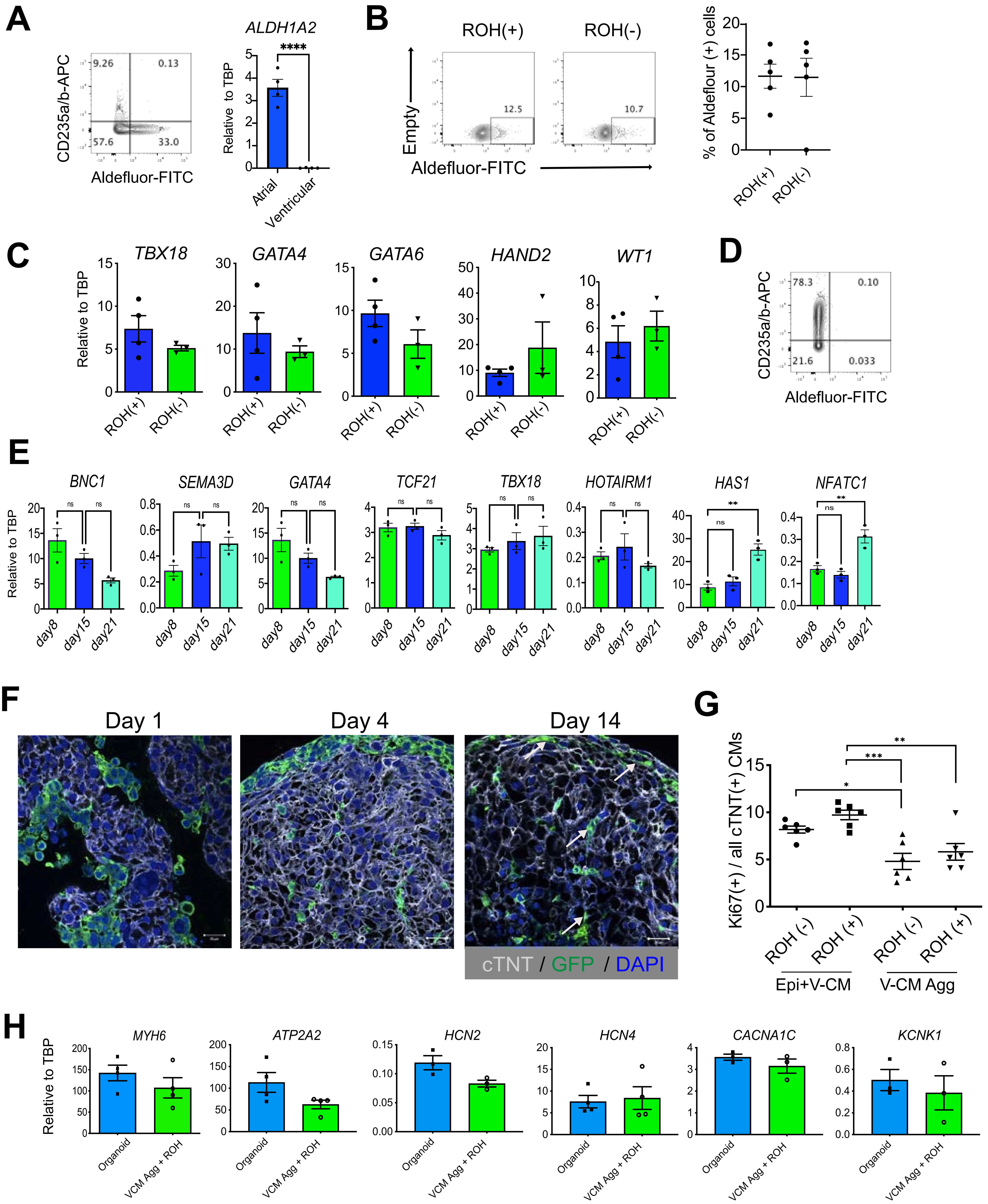

**Supplementary Figure 1. Generation of cardiac organoids with ventricular cardiomyocytes and epicardial cells.** (A) Left; Representative flow cytometry analyses of Aldefluor and CD235a/b in day 4 atrial mesoderm. Right; RT-qPCR expression analyses of *ALDH1A2* in atrial and ventricular mesoderm (N=4). (B) Left; Representative flow cytometry analysis of Aldefluor in ROH-treated and untreated epicardial cells at day 8. Right; Quantification of the percentage of Aldefluor-positive cells in the indicated conditions (N=5). (C) RT-qPCR expression analyses of epicardial-related and cardiac transcription factors (*TBX18, GATA4, GATA6, HAND2, WT1*) in the ROH-treated and untreated (ROH (-)) epicardial cells (N=3-4). (D) Representative flow cytometry analyses of Aldefluor and CD235a/b in day 4 ventricular mesoderm. (E) RT-qPCR expression analyses of epicardial-related genes (*BNC1, TBX18, GATA4, SEMA3D, TCF21, HOTAIRM1, HAS1, NFATC1*) in the epicardial cells at the indicated time of culture (N=3). (F) Representative immunostaining of GFP and cTNT in day1, 4, 14 cardiac organoids. White arrows indicate elongated GFP-positive EPDCs. GFP; epi-derived cells (EPDCs), Scale bar; 20um. (G) Quantification of the changes in the percentage of Ki67-positive cardiomyocytes in the cardiac organoids and cardiomyocyte aggregates (VCM Agg) treated as indicated (N=6). (H) RT-qPCR expression analyses of sarcomere-related (*MYH6*), $Ca^{2+}$ handling-related (*ATP2A2*) and ion channel-related genes (*HCN2, HCN4, CACNA1C, KCNK1*) in the indicated conditions (N=3-4). *p<0.05, **p<0.01, ***p<0.001 by unpaired t-test (A) and by one-way ANOVA with Tukey's multiple comparisons (E) and (G). All error bars represent SEM.

### Supplementary Figure 2.

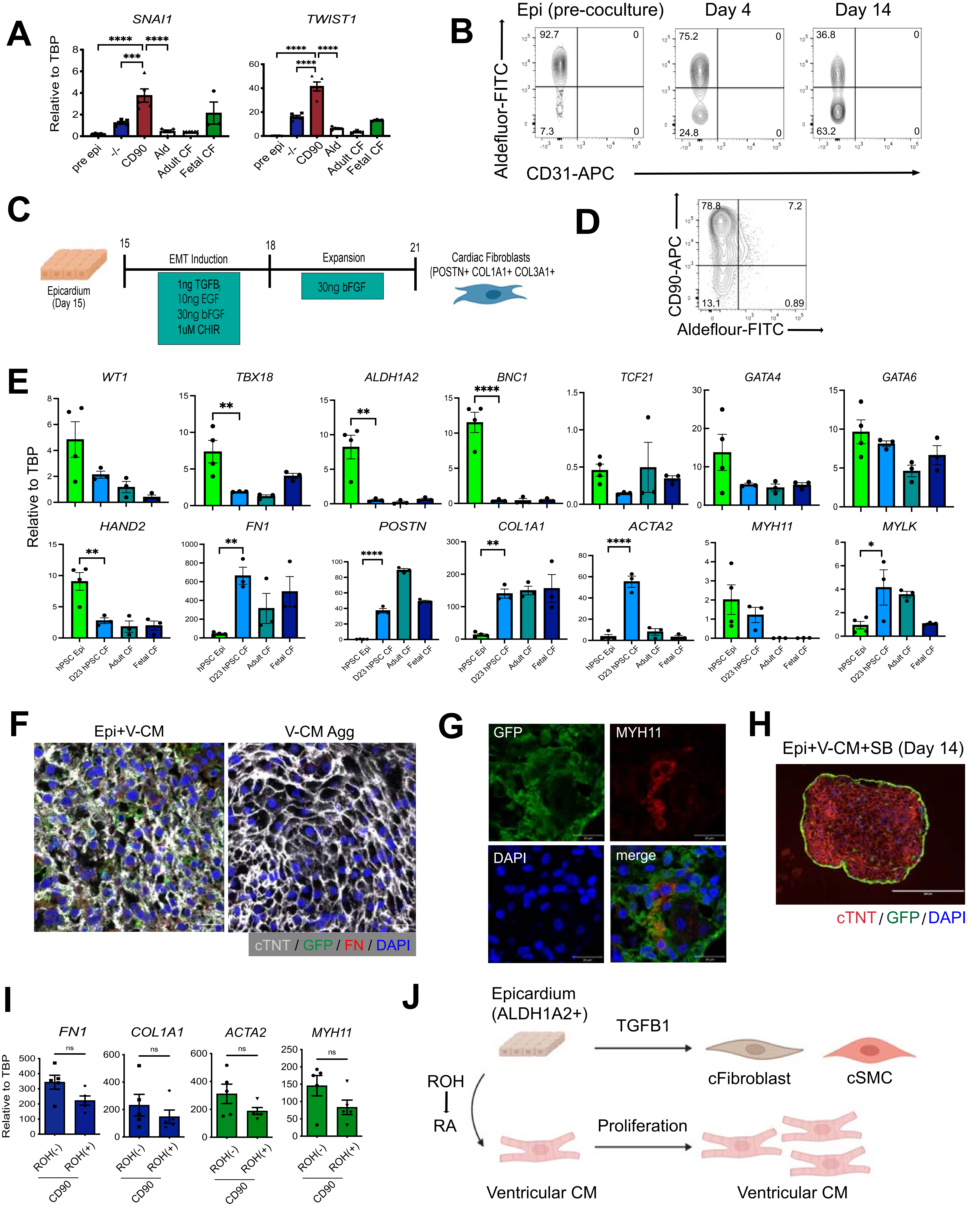

**Supplementary Figure 2. Characterization epicardial-derived cells (EPDCs) in the cardiac organoids.** (A) RT-qPCR expression analyses of EMT-related (*SNAI1*, *TWIST1*) genes in the indicated populations (N=4-5). Adult and fetal CF was included as a reference (N=3 each). Epi: day15 epicardium prior to coculture. (B) Representative flow cytometry analysis of Aldefluor and CD31 in EPDCs in the cardiac organoids in the indicated time points. (C) Protocol to directly differentiate hPSC-derived epicardial cells into cardiac fibroblasts. (D) Representative flow cytometry analysis of CD90 and Aldefluor in directly differentiated CFs derived from epicardial cells. (E) RT-qPCR expression analyses of epicardial (*WT1, TBX18, ALDH1A2*, *BNC1*), transcription factor (*TCF21, GATA4, GATA6, HAND2)*, extracellular matrix (*FN1, POSTN, COL1A1*), and smooth muscle (*ACTA2, MYH11, MYLK*) genes in the indicated populations (N=3). Adult and fetal CF were included as a reference (N=3 each). Epi: day15 epicardium, (F) Representative immunostaining of fibronectin (FN) and GFP (EPDCs) in the cardiac organoids and ventricular cardiomyocyte aggregates (V-CM Agg). Scale bar; 20um. (G) Representative immunostaining of MYH11 in the GFP-positive EPDCs in the cardiac organoids. Scale bar; 20um. (H) Representative immunostaining of day 14 SB-treated cardiac organoids. Scale bar; 200um. (I) RT-qPCR expression analyses of *FN1*, *COL1, ACTA2,* and *MYH11* in CD90-positive populations in the ROH-treated and untreated organoids (N=5). (J) Schematic of the cardiac organoids showing the interaction between the cardiomyocytes and the epicardial cells. ****$p<0.0001$, *** $p<0.001$, ** $p<0.01$, *$p<0.05$, by one-way ANOVA with Tukey's multiple comparisons in (A) and by unpaired t test in (E)(I).

### Supplementary Figure 3.

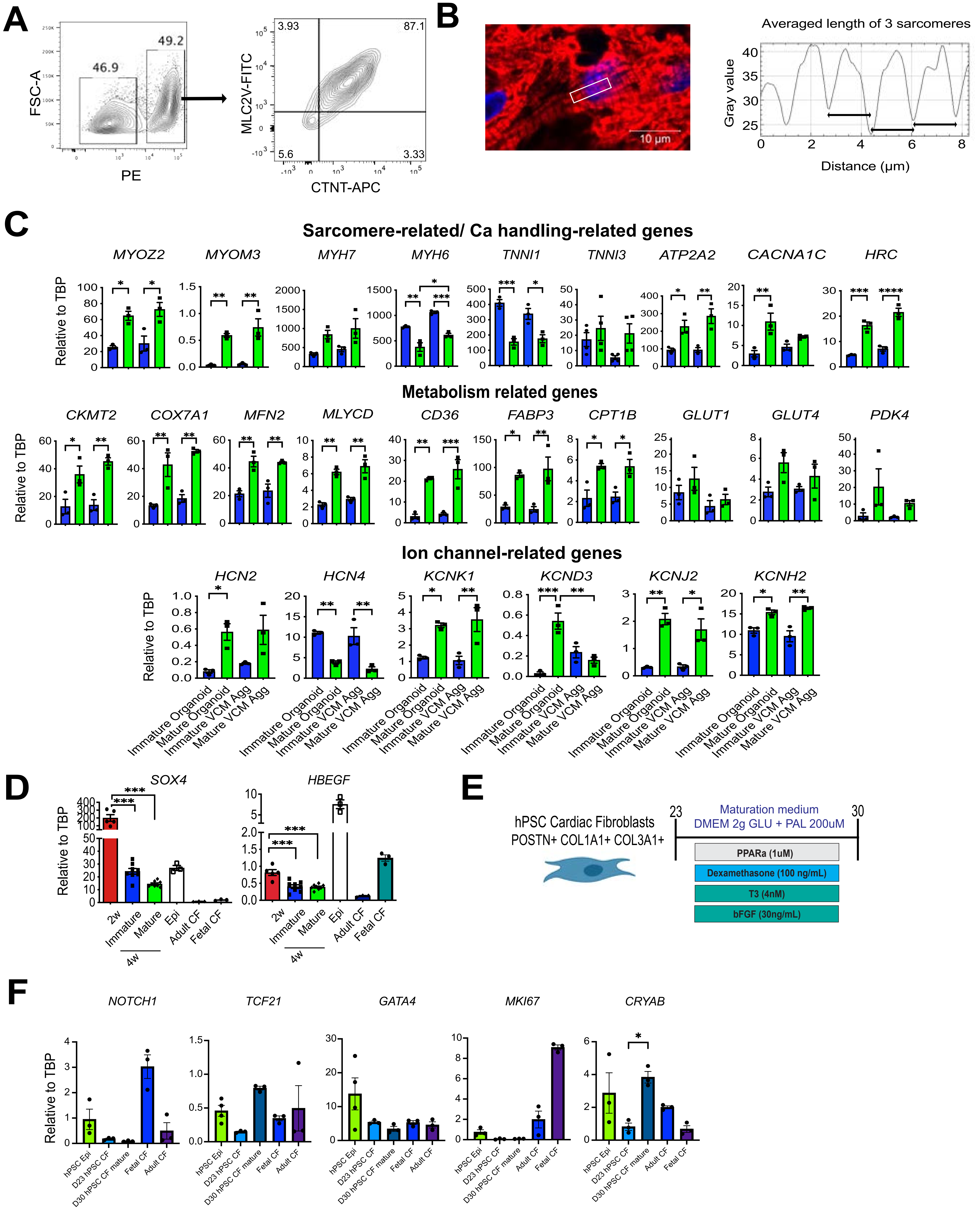

**Supplementary Figure 3. Metabolic maturation in cardiac organoids.** (A) Representative flow cytometry analysis of cTNT and MLC2v in $RFP^+$ sorted population in the cardiac organoids. (B) The method to measure sarcomere length in an image of cTNT staining. Sarcomere length was measured in an area with a white square by Image J. Scale bar; 10um. (C) RT-qPCR expression analyses of the sarcomere (*MYOZ2, MYOM3, DES, MYH7, MYH6, TNNI1, TNNI3*), Ca handling (*ATP2A2, HRC*), fatty acid oxidation (*CD36, FABP3, CPT1B, MLYCD*) and mitochondria (*CKMT2, COX7A1, MFN2*), ion channel (*KCNK1, HCN2, KCND3, CACNA1C, KCNJ2, KCNH2, HCN4*), glycolysis (*GLUT1, GLUT4, PDK4*) -related genes in FACS isolated cardiomyocyte from the immature and mature cardiac organoids and immature and mature ventricular cardiomyocyte aggregates (VCM Agg) (N=3). (D) RT-qPCR expression analyses of *SOX4* and *HBEGF* in FACS isolated $CD90^+$ cardiac fibroblasts populations from the immature and mature cardiac organoids (N=4-8). Adult and fetal CF was included as a reference (N=3 each). epi: day15 epicardial cells prior to the initiation of coculture (N=3, 4), CF: cardiac fibroblasts. (E) Protocol to culture directly differentiate hPSC-derived epicardial cells in the maturation media. (F) RT-qPCR expression analyses of indicated genes in the day23 CFs and day30 CFs cultured in the maturation media (N=3). Adult and fetal CF were included as a reference (N=3). epi: day15 epicardial cells (N=3, 4), *p<0.05, **p<0.01, ***p<0.001, ****p<0.0001 by one-way ANOVA with Tukey's multiple comparisons in (C) (D) (F). All error bars represent SEM.

### Supplementary Figure 4.

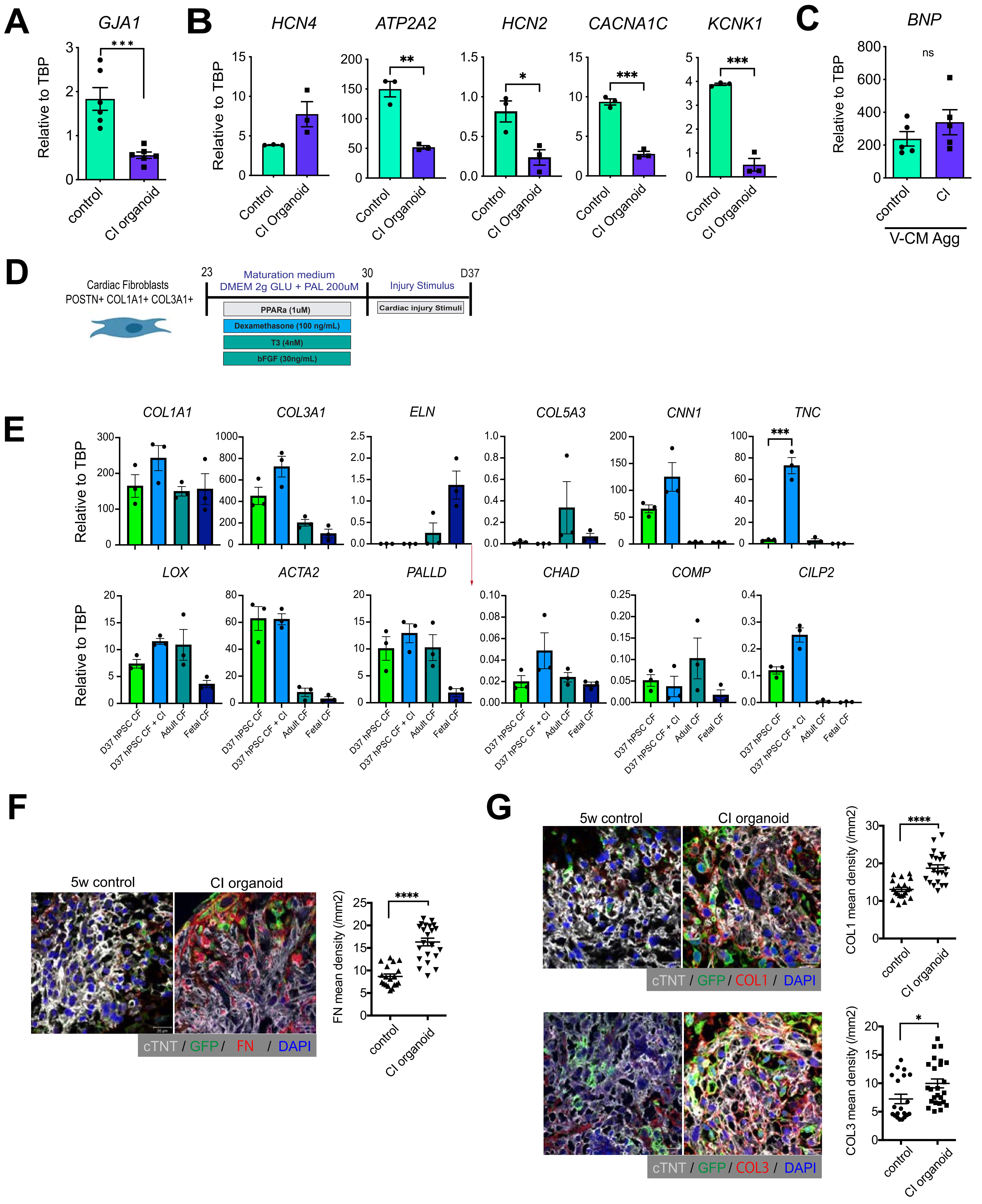

**Supplementary Figure 4. Modeling cardiac injury in cardiac organoids.** (A) RT-qPCR expression analyses of *GJA1* in the FACS isolated cardiomyocyte populations from the control and CI organoids (N=6). (B) RT-qPCR expression analyses of $Ca^{2+}$ handling-related (*ATP2A2*) and ion channel-related (*HCN4, HCN2, CACNA1C, KCNK1*) in the FACS isolated cardiomyocyte populations from the control and CI organoids (N=3). (C) RT-qPCR expression analyses of heart failure marker (*BNP*) in the ventricular cardiomyocyte aggregates (V-CM-Agg) cultured as indicated (N=5). (D) Protocol to expose the cardiac injury stimuli in directly differentiated CFs derived from hPSC-epicardial cells. (E) RT-qPCR expression analyses of extracellular matrix (*COL1A1, COL3A1, ELN, COL5A3*), cytoskeleton (*ACTA2, PALLD, CNN1*), fibrosis-related (*TNC, LOX*), and matrifibrocyte-related (*CHAD, COMP, CILP2*) genes in the indicated populations (N=3 for matrifibrocyte genes, N=6 for others). Adult and fetal CF was included as a reference (N=3 each). (F) Left; Representative immunostaining of fibronectin (FN) in the control and CI organoids at 5 weeks of culture. Scale bar; 20um. Right; Quantification of the FN density in the two organoid populations (N=19-22). (G) Upper left; Representative immunostaining of collagen type 1 (COL1) in the control and CI organoids at 5 weeks of culture. Scale bar; 20um. Upper right; Quantification of the COL1 density in the two organoid populations (N=20-21). Lower left; Representative immunostaining of collagen type 3 (COL3) in the control and CI organoids at 5 weeks of culture. Scale bar; 20um. Lower right; Quantification of the COL3 density in two organoid populations (N=20-25). CI organoid: cardiac injury organoid. CF: cardiac fibroblasts. *p<0.05, **p<0.01, ***p<0.001, ****p<0.0001 by unpaired t-test. All error bars represent SEM.

### Supplementary Figure 5.

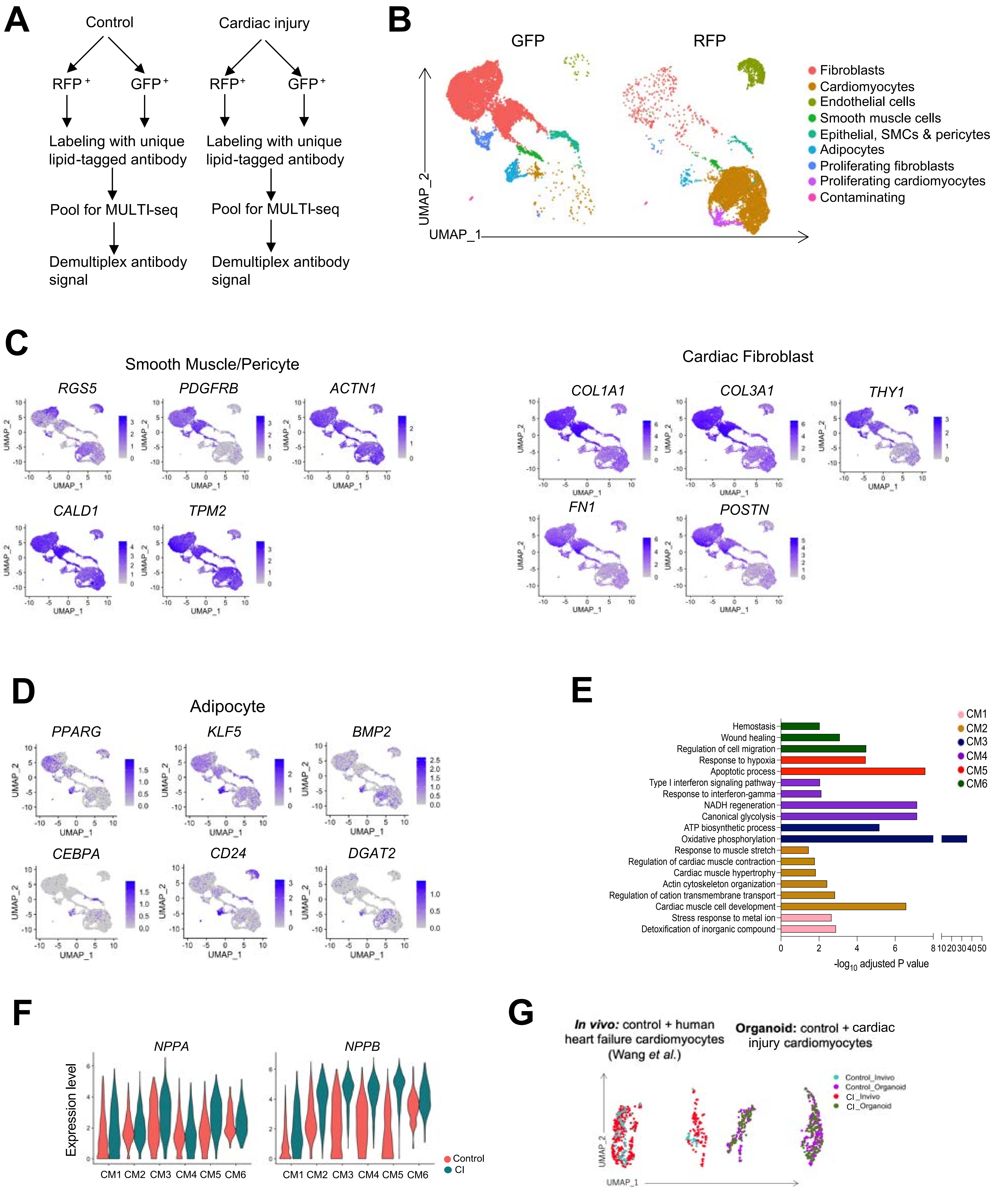

**Supplementary Figure 5. Single-cell RNA sequencing analyses of the cardiac organoids.** (A) Flow chart of single-cell RNA sequencing sample workflow. RFP-positive and GFP-positive fractions of control and CI organoids were sorted and labelled individually using lipid-tagged antibodies, and subsequently pooled for sequencing using the 10x Genomics platform and MULTI-seq. Following sequencing, the data was demultiplexed to assign cells back to RFP or GFP fractions. (B) UMAP dimensionality reduction of total cells in control and CI samples split by RFP and GFP fractions. (C) UMAP plots showing expression of smooth muscle/pericyte and fibroblast-related genes (*RGS5, PDGFRB, ACTN1, CALD1, TPM2, COL1A1, COL3A1, FN1, POSTN, THY1*). (D) UMAP plots showing expression of adipocyte-related genes (*PPARG, KLF5, BMP2, CEBPA, CD24, DGAT2*). (E) Pathway enrichment analysis was performed for each cardiomyocyte cluster in Figure 5 (gProfiler, GO: Biological Processes). The top unique pathways in each cluster are displayed with their enrichment score ($-\log_{10}$ of adjusted p-value). (F) Violin plots showing expression of *NPPA* and *NPPB* in each cardiomyocyte cluster in control and CI samples. (G) Integration analysis of cardiomyocyte population in control and CI organoid samples with cardiomyocytes from *in vivo* human coronary heart failure dataset (Wang *et al.*). Datasets were downsampled to 400 cells each to avoid biases associated with differential cell numbers. CM: cardiomyocyte, CI organoid: cardiac injury organoid.

### Supplementary Figure 6.

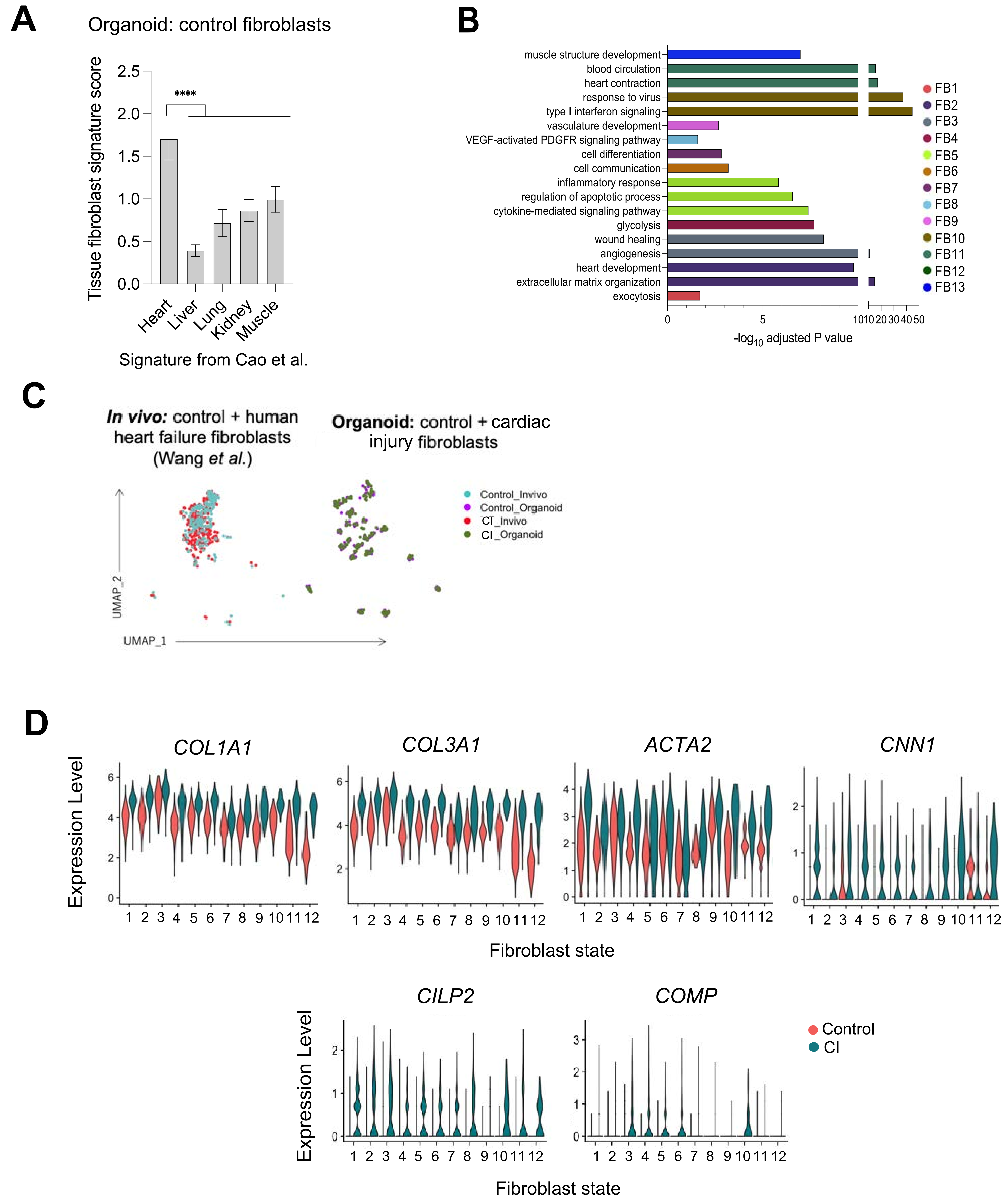

**Supplementary Figure 6. Molecular characterization of fibroblasts in the cardiac organoids.** (A) Single-cell RNA sequence data of human fetal fibroblasts from 5 tissues (Cao *et al.*) were compared to identify tissue-specific fibroblast signatures. Each tissue-specific fibroblast signature was used to score control organoid fibroblasts. One-way ANOVA was performed for each tissue signature compared to the heart signature. ****$p<0.0001$. (B) Pathway enrichment analysis was performed for each fibroblast cluster in Figure 6 (gProfiler, GO: Biological Processes). The top unique pathways in each cluster are displayed with their enrichment score ($-\log_{10}$ of adjusted p-value). (C) Integration analysis of fibroblast population in control and CI organoid samples with fibroblasts from *in vivo* human coronary heart failure dataset (Wang *et al.*). Datasets were downsampled to 400 cells each to avoid biases associated with differential cell numbers. (D) Violin plots showing expression of fibrosis-related genes (*COL1A1, COL3A1, ACTA2, CNN1, CILP2, COMP*) in each fibroblast cluster in control and CI samples. FB: fibroblast, CI organoid: cardiac injury organoid.

### Supplementary Figure 7.

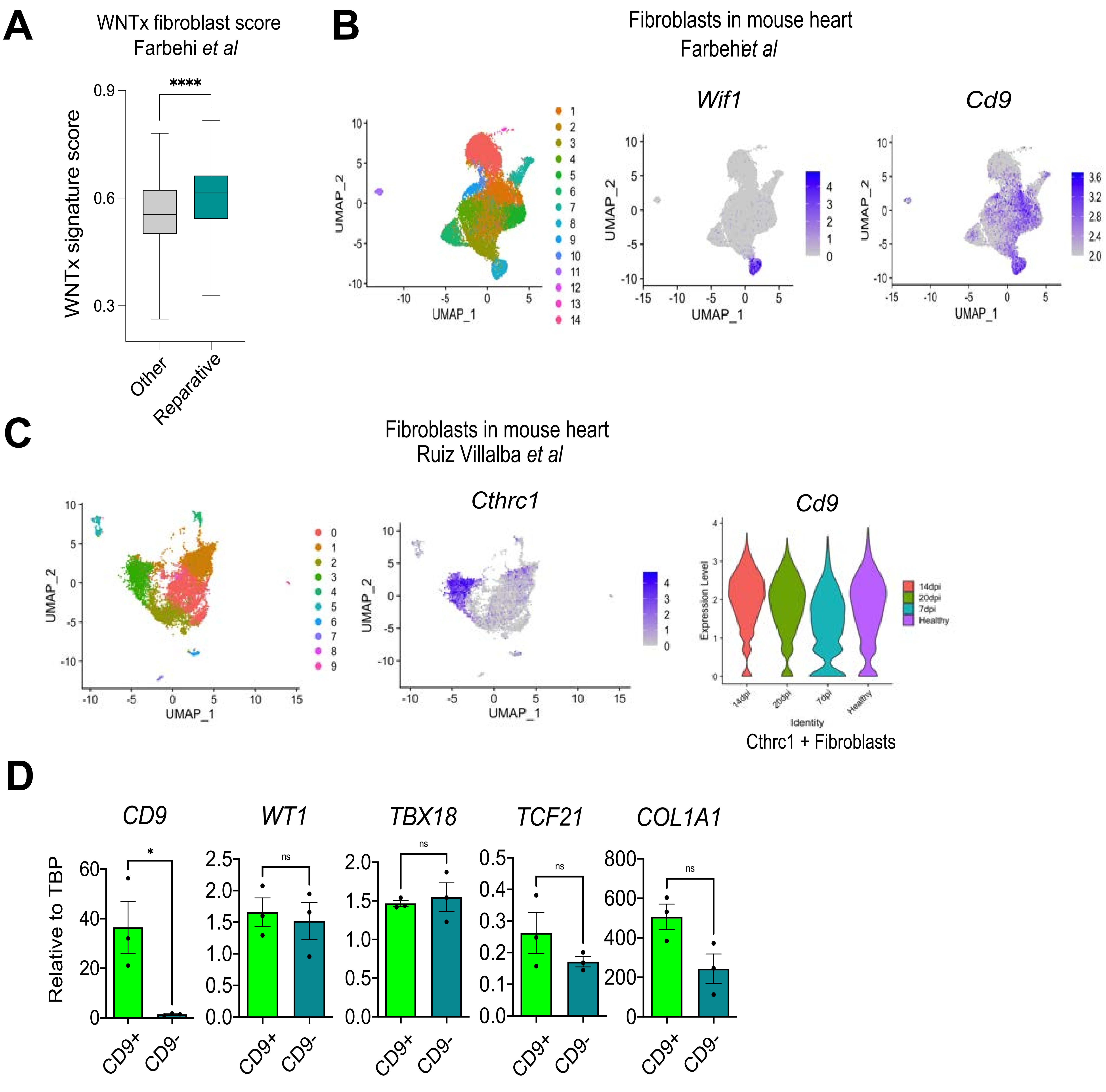

**Supplementary Figure 7. Identification of a CD9$^+$ reparative like fibroblast population** (A) A signature for the WNTx fibroblast cluster defined by Farbehi *et al*. was generated by comparing the WTNx cluster to all other fibroblasts. “Reparative” and “other” fibroblasts from the CI organoids (see Figure 7) were scored with this signature. An unpaired t-test was performed to compare the two groups. (B) Clustering analysis of fibroblast populations and UMAP plots showing expression of *Wif1* and *Cd9* in mouse heart *in vivo* (Farberi *et al*.) (C) Left; Clustering analysis of fibroblast populations and UMAP plots showing expression of *Cthrc1*. Right; Violin plots showing expression of *Cd9* in Cthrc1-positive fibroblasts at the indicated time points. All data are analyzed from the mouse heart dataset of Ruiz Villalba *et al*. (D) RT-qPCR expression analyses of indicated genes in the CD9$^+$ and CD9$^-$ cardiac fibroblast populations isolated from the CI organoids (N=3). CI organoid: cardiac injury organoid. ****$p<0.0001$, *** $p<0.001$,** $p<0.01$, *$p<0.05$, by unpaired t-test.

### Supplementary Figure 8.

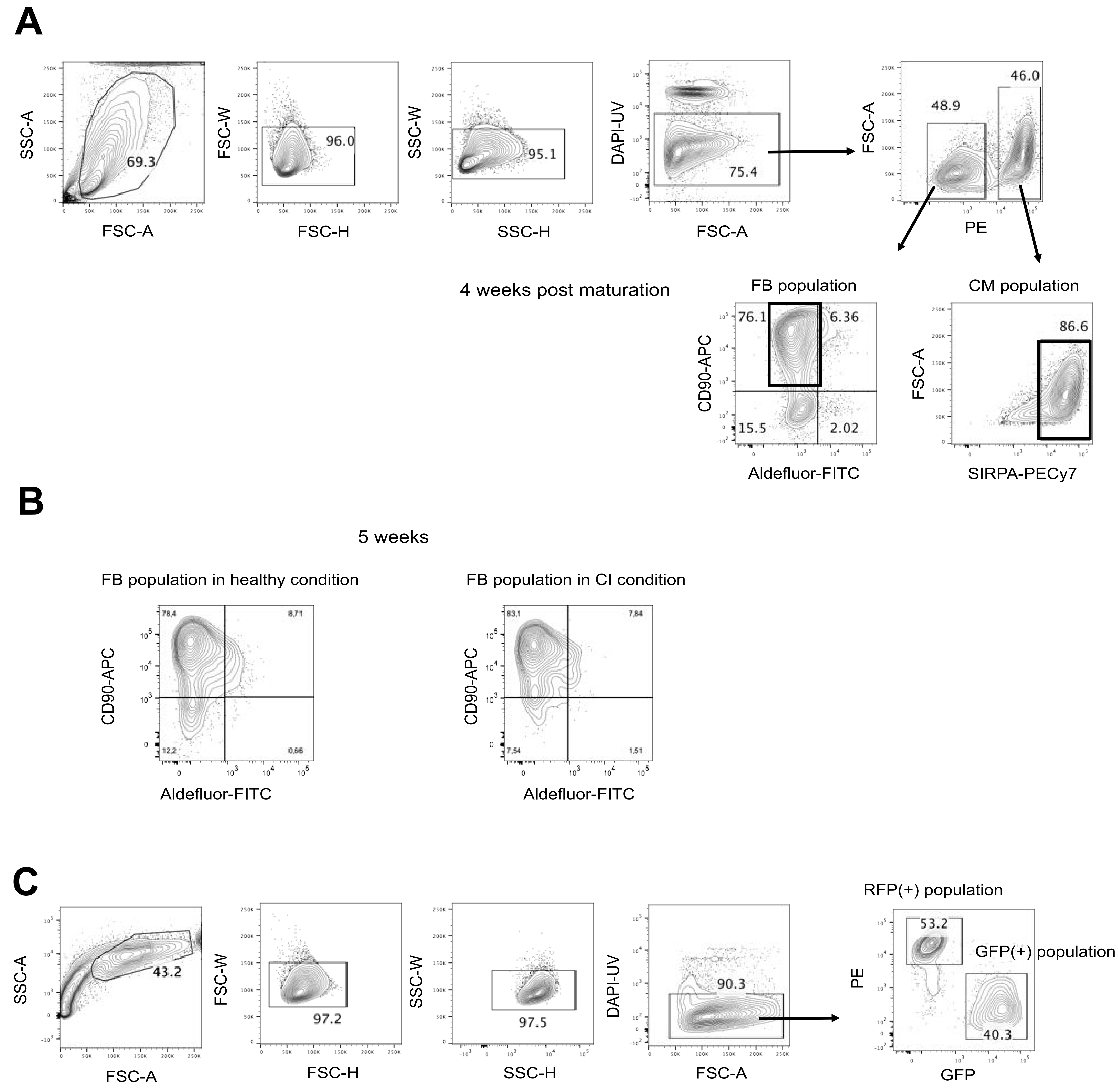

**Supplementary Figure 8. Gating strategy for flow-cytometric analysis.** (A) Gating strategy for sorting RFP(+) / SIRPA(+)-cardiomyocytes and CD90(+)-fibroblasts derived from epicardial cells in the cardiac organoid used in Figure 2-4 and Supplementary Figure 2-4. Representative flow cytometry result of CD90 and Aldefluor in EPDC population in the 4-week cardiac organoids are shown. (B) Representative flow cytometry analysis of CD90 and Aldefluor in EPDC population in the 5-week cardiac organoids with or without cardiac injury. (C) Gating strategy for sorting RFP-positive and GFP-positive fractions of the cardiac organoids for MULTI-seq shown in Figure 5, 6 and Supplementary Figure 5, 6.
